# Multiple 9-1-1 complexes promote homolog synapsis, DSB repair, and ATR signaling during mammalian meiosis

**DOI:** 10.1101/2021.04.09.439198

**Authors:** Catalina Pereira, Gerardo A. Arroyo-Martinez, Matthew Z. Guo, Michael S. Downey, Emma R. Kelly, Kathryn J. Grive, Shantha K. Mahadevaiah, Jennie Sims, Vitor Marcel Faça, Charlton Tsai, Carl J. Schiltz, Niek Wit, Heinz Jacobs, Nathan L. Clark, Raimundo Freire, James M. A. Turner, Amy M. Lyndaker, Miguel A. Brieño-Enríquez, Paula E. Cohen, Marcus B. Smolka, Robert S. Weiss

## Abstract

DNA damage response mechanisms have meiotic roles that ensure successful gamete formation. While completion of meiotic double-strand break (DSB) repair requires the canonical RAD9A-RAD1-HUS1 (9A-1-1) complex, mammalian meiocytes also express RAD9A and HUS1 paralogs, RAD9B and HUS1B, predicted to form alternative 9-1-1 complexes. The RAD1 subunit is shared by all predicted 9-1-1 complexes and localizes to meiotic chromosomes even in the absence of HUS1 and RAD9A. Here we report that testis-specific RAD1 disruption resulted in impaired DSB repair, germ cell depletion and infertility. Unlike *Hus1* or *Rad9a* disruption, *Rad1* loss also caused defects in homolog synapsis, ATR signaling and meiotic sex chromosome inactivation. Comprehensive testis phosphoproteomics revealed that RAD1 and ATR coordinately regulate numerous proteins involved in DSB repair, meiotic silencing, synaptonemal complex formation, and cohesion. Together, these results establish critical roles for both canonical and alternative 9-1-1 complexes in meiotic ATR activation and successful prophase I completion.

## INTRODUCTION

DNA damage response (DDR) mechanisms protect genomic integrity by sensing and repairing DNA lesions or initiating apoptosis when lesions are unrepairable (Blackford and Jackson, 2017). DDR proteins are also essential for proper haploid gamete formation. Although double-strand DNA breaks (DSBs) are considered to be the most toxic form of DNA damage, meiotic recombination relies on SPO11-induced DSBs for homologous chromosomes to synapse, exchange genetic material, and properly segregate at the first meiotic division (Bolcun-Filas et al., 2014; Gray and Cohen, 2016). Of particular importance are the meiotic events that occur during the five sub-stages of prophase I, a major feature of which involves the transient formation of the proteinaceous structure called the synaptonemal complex (SC) (Cahoon and Hawley, 2016; Gray and Cohen, 2016). During the first stage, leptonema, axial elements containing SC protein 3 (SYCP3) form along condensed chromosomes (Page and Hawley, 2004). Additionally, the DNA damage marker, γH2AX, accumulates during leptonema as chromosomes experience SPO11-induced DSBs. Progression into zygonema is characterized by the pairing and synapsis of chromosomes, marked by the presence of the central element protein SC protein 1 (SYCP1). By pachynema DSB repair is completed and γH2AX is no longer present on the fully synapsed autosomes. However, in male meiocytes, abundant γH2AX is apparent at the sex body containing the X and Y chromosomes, which synapse only in a small domain called the pseudoautosomal region but otherwise remain unsynapsed. Meiotic cells subsequently enter diplonema, featuring dissolution of the central element while homologous chromosomes remain tethered by crossovers. Breakdown of the SC marks the final stage in prophase I, diakinesis.

Ataxia-telangiectasia and Rad3-related (ATR) kinase is a key regulator of recombinational DSB repair and synapsis throughout meiotic prophase I (Pereira et al., 2020). ATR activation in somatic cells has been well characterized; however, the mechanisms of meiotic ATR activation have not been fully elucidated. ATR activation in response to replication stress and other signals in mitotic cells is known to involve interaction between the RAD9A-RAD1-HUS1 (9A-1-1) complex and Topoisomerase 2-Binding Protein I (TOPBP1) (Blackford and Jackson, 2017). The toroidal, PCNA-like 9A-1-1 complex is loaded at recessed DNA ends by the RAD17-Replication Factor C (RFC) clamp loader (Eichinger and Jentsch, 2011). ATR in association with ATR Interacting Protein (ATRIP) independently localizes to Replication Protein A (RPA)-coated single stranded DNA (Zou, 2003) The 9A-1-1 complex then interacts with the RAD9A-RAD1-HUS1 Interacting Nuclear Orphan (RHINO) and TOPBP1, which allows TOPBP1 to activate ATR via its ATR-activating domain (Cotta-Ramusino et al., 2011; Delacroix et al., 2007; Lindsey-Boltz et al., 2015) ATR activation initiates several downstream processes such as cell cycle arrest, DNA repair, fork stabilization, and inhibition of new origin firing, or triggers apoptosis (Saldivar et al., 2017). Independent of 9A-1-1/TOPBP1, ATR also can be directly activated during a normal mitotic cell cycle by Ewing’s Tumor Associated-antigen 1 (ETAA1), in part to promote metaphase chromosome alignment and spindle assembly checkpoint function (Bass and Cortez, 2019). During meiotic prophase I, homologous chromosomes pair and undergo recombination, with regions of asynapsis being subjected to DDR-dependent transcriptional silencing. ATR, along with meiosis specific HORMA (Hop1, Rev7, and Mad2)-domain proteins, TOPBP1, and other factors, localizes to unsynapsed chromatin regions in leptotene-and zygotene-stage cells (Fedoriw et al., 2015). At pachynema, the homologs are fully synapsed, at which point ATR localizes only to the unsynapsed axes and throughout the chromatin of the X and Y chromosomes, where it triggers a mechanism called meiotic sex chromosome inactivation (MSCI). MSCI is essential for successful meiotic progression through the silencing of toxic Y-linked genes and sequestration of DDR proteins away from autosomes (Abe et al., 2020; Royo et al., 2010; Turner, 2015). Central to MSCI is ATR-mediated phosphorylation of BRCA1 and H2AX on chromatin loops (Fukuda et al., 2012; Royo et al., 2013; Turner et al., 2004). Similarly, ATR mediates meiotic silencing of unsynapsed chromatin (MSUC) at autosomes that have failed to synapse properly (Turner, 2007, 2015). Beyond silencing, ATR has an essential role in promoting RAD51 and DMC1 loading to enable meiotic DSB repair (Pacheco et al., 2018; Widger et al., 2018). Previous work indicates that HUS1 and RAD9A are largely dispensable for meiotic ATR activation (Lyndaker et al., 2013a; Vasileva et al., 2013), raising the intriguing possibility that HUS1B-and RAD9B-containing alternative 9-1-1 complexes contribute to ATR activation during mammalian meiosis.

In addition to its checkpoint activation role, the 9A-1-1 complex also functions as a molecular scaffold for proteins in multiple DNA repair pathways. For example, the 9A-1-1 complex participates in homologous recombination by interacting with the RAD51 recombinase (Pandita et al., 2006) and EXO1 exonuclease (Karras et al., 2013; Ngo et al., 2014; Ngo and Lydall, 2015). Consistent with these observations from mitotic cells, RAD9A co-localizes with RAD51 on meiotic chromosome cores (Lyndaker et al., 2013a). In wild-type pachytene-stage cells, RAD51 foci are lost as DSBs are resolved, whereas without *Hus1* RAD51 is retained on spermatocyte autosomes into late prophase I (Lyndaker et al., 2013a).

Loss of any canonical 9A-1-1 subunit in mice leads to embryonic lethality (Han et al., 2010; Hopkins et al., 2004; Weiss et al., 2000). In conditional knockout (CKO) models, loss of *Hus1* or *Rad9a* in the testis results in persistent DSBs during meiotic prophase leading to reduced testis size, decreased sperm count, and sub-fertility (Lyndaker et al., 2013a; Vasileva et al., 2013). Interestingly, localization of RAD1 and RAD9A to meiotic chromosome cores only partially overlaps, with RAD1 localizing to asynapsed chromosomes and along the entire X chromosome, and RAD9A in a more punctate pattern suggestive of DSB sites (Freire et al., 1998; Lyndaker et al., 2013a). Although RAD9A fails to localize properly in *Hus1*-deficient meiocytes, RAD1 localization to meiotic chromosome cores is largely HUS1-independent, supporting the idea that RAD1 can act outside of the canonical 9A-1-1 complex.

The HUS1 and RAD9A paralogs, HUS1B and RAD9B, are highly expressed in testis (Hang et al., 2002; Hopkins et al., 2004). Based on our previous results and the findings discussed above, we previously hypothesized that meiocytes contain alternative 9-1-1 complexes, RAD9B-RAD1-HUS1 (9B-1-1) and RAD9B-RAD1-HUS1B (9B-1-1B) (Lyndaker et al., 2013b). Since RAD1 has no known paralogs, it is expected to be common to both canonical and alternative 9-1-1 complexes. In order to elucidate the roles of each of the 9-1-1 complexes in mammalian meiosis, we generated *Rad1* CKO mice in which *Rad1* was disrupted specifically in male spermatocytes. *Rad1* CKO mice exhibited reduced sperm count, reduced testis size and severe germ cell loss associated with DSB repair defects, consistent with previous studies of HUS1 and RAD9A. However, homolog synapsis and MSCI, which were largely unaffected in *Hus1* or *Rad9a* CKO mice, were disrupted by *Rad1* loss. Whole testis-phosphoproteomic analyses further highlighted the importance of multiple 9-1-1 complexes in ATR-mediated processes such as meiotic silencing and cohesin regulation. This study highlights the importance of canonical and alternative 9-1-1 complexes during mammalian meiosis and establishes key roles for these DDR clamps in ATR activation, homolog synapsis and MSCI.

## RESULTS

### Evolution and tissue-specific expression of 9-1-1 subunits

Human RAD9A and RAD9B share 45% identity, while HUS1 and HUS1B are 48% identical (Dufault et al., 2003; Hang et al., 2002). Inspection of genomic sequences revealed that *Rad9b* genes are present in the syntenic genomic region of all placental species analyzed, whereas *Rad9a* was likely lost in a few species, including wallaby, tree shrew and sloth (Figure 1A-1B). Phylogenetic analysis suggested that the duplication event generating *Rad9a* and *Rad9b* occurred prior to the evolution of bony fish ancestors (*D. rerio*), whereas the single exon *Hus1b* gene likely arose after a retrocopy duplication event later in evolution in mammals. Ortholog matrix and evolutionary tree analyses of placental mammals further showed that *Rad1* is highly conserved, with no identifiable paralog.

**Figure 1:**
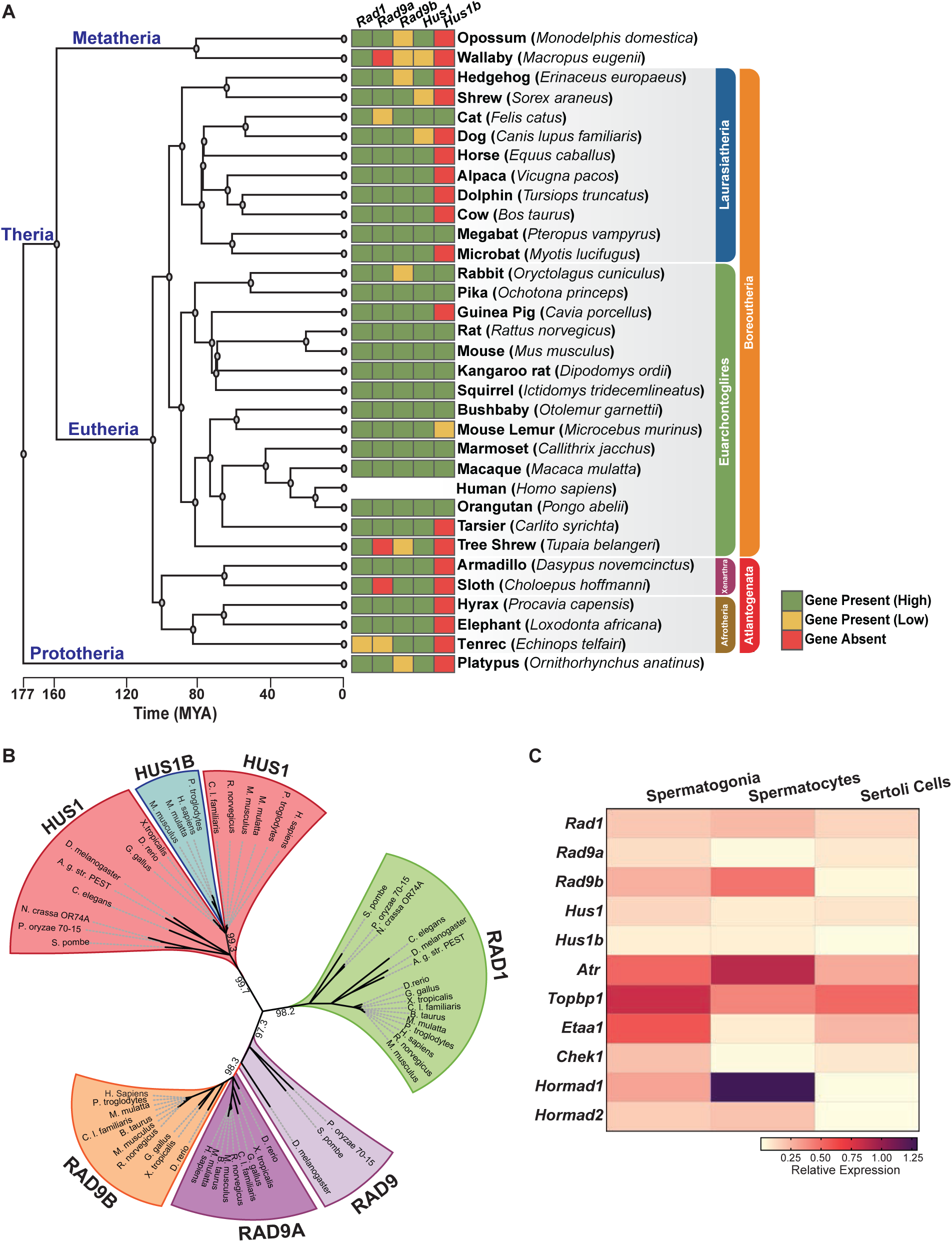
Phylogenetic analysis of 9-1-1 complex subunits. (A) Gene presence and absence matrix of human 9-1-1 subunit orthologue genes in 33 representative mammals. High confidence was determined if the genomic sequence had ≥50% of both target and query sequence identity, and a pairwise whole genomic alignment score of ≥50 when compared to human or if the genomic region containing the gene was syntenic with human. If an ortholog did not reach the threshold then it was annotated as low confidence (yellow). If no ortholog was found, then it was considered absent (red). A Cladogram was obtained from timetree.org. (B) Maximum likelihood unrooted phylogenetic tree of 9-1-1 subunit genes based on JTT+I+G+F. Protein sequences were obtained from NCBI HomoloGene and include: bacteria (*Pleomorphomonas oryzae*), fungi (*Schizosaccharomyces pombe, Neurospora crassa*), nematode (*Caenorhabditis elegans)*, true flies (*Drosophila melanogaster, Anopheles gambiae str. Pest*), fish (*Danio rerio*), frog (*Xenopus tropicalis*), bird (*Gallus gallus)*, carnivora (*Canis lupus*), rodents (*Rattus norvegicus, Mus musculus*) and primates (*Homo sapiens, Mus musculus, Macaca mulatta, Pan troglodytes*). Sequences were aligned by Clustal Omega and substitution model was tested on ProtTest. Ultrafast bootstrap (x1000 replicates) was performed in IQ-TREE web server and nodes below 70% branch support were collapsed. Branch distance represents substitution rate. (C) Heatmap of single cell RNA sequencing data from mouse testes was queried to assess the expression of 9-1-1 subunits, *Atr, Topbp1*, *Etaa1*, *Chk1*, *Hormad1* and *Hormad2* in spermatogonia, spermatocytes and Sertoli cells. Expression of *Rad9b* in spermatocytes p value ≤ 5.47e^-10^, *Hus1b* p value ≤1.20e^-09^, *Rad1* p value ≤1.08e^-08^. Expression of *Rad9a* and *Hus1* in spermatogonia p value ≤5.61e^-19^; p value ≤3.78e^-09^. Relative expression is shown of each gene, highest expression observed in purple and lowest expression observed in yellow.

Human and mouse gene expression data indicate that the 9-1-1 paralogs are highly expressed in testes but no other tissues, hinting at a potential role for RAD9B and HUS1B in spermatogenesis (Figure 1-figure supplement 1A). To further define the cell type-specific expression patterns of the 9-1-1 subunits within the testes, we mined single-cell RNA sequencing data from wild-type adult mouse testis (Grive et al., 2019), comparing relative expression in spermatogonia, spermatocytes, and Sertoli cells. *Rad9b* expression was highest in spermatocytes as compared to spermatogonia and Sertoli cells (Figure 1C). Similarly, *Rad1* and *Hus1b* expression were highest in spermatocytes (Figure 1C). Conversely, *Rad9a* and *Hus1* relative gene expression was highest in spermatogonia (Figure 1C). As expected, we found expression of *Atr* and meiotic silencing genes *Hormad1* and *Hormad2* to be significantly higher in spermatocytes than spermatogonia or Sertoli cells. Spermatogonia also displayed relatively high levels of *Atr*, along with *Topbp1* and *Etaa1* (Figure 1C). Analysis of expression data from human testis similarly showed that *Hus1b* and *Rad9b* expression was highest in spermatocytes, whereas *Rad1* and *Rad9a* expression was highest in spermatogonia and *Hus1* expression was highest in early spermatids (Guo et al., 2018). Together these results suggest that the 9-1-1/TOPBP1/ATR and ETAA1/ATR signaling axes are expressed in pre-meiotic spermatogonia and suggest roles for alternative 9-1-1 complexes in male meiosis.

To further analyze the evolutionary relationships between 9-1-1 subunits, we performed evolutionary rate covariation (ERC) analysis, which assesses correlations in gene evolutionary history and can reveal functionally significant relationships (Clark et al., 2012; Wolfe and Clark, 2015). ERC analysis was performed between all of the 9-1-1 subunits in a pairwise fashion across 33 mammalian species (Clark et al., 2012). Significant ERC values were identified between the RAD1, HUS1, and RAD9B subunits, supporting the notion that alternative 9-1-1 complexes assemble in germ cells (Figure 2A). These findings are consistent with reports that RAD9B physically interacts with RAD1, HUS1, and HUS1B (Dufault et al., 2003), and similarly that HUS1B interacts with RAD1 (Hang et al., 2002), suggesting that the paralogs contribute to alternative 9-1-1 complexes that include RAD9B-RAD1-HUS1 (9B-1-1) and RAD9B-HUS1-HUS1B (9B-1-1B) (Figure 2B).

**Figure 2:**
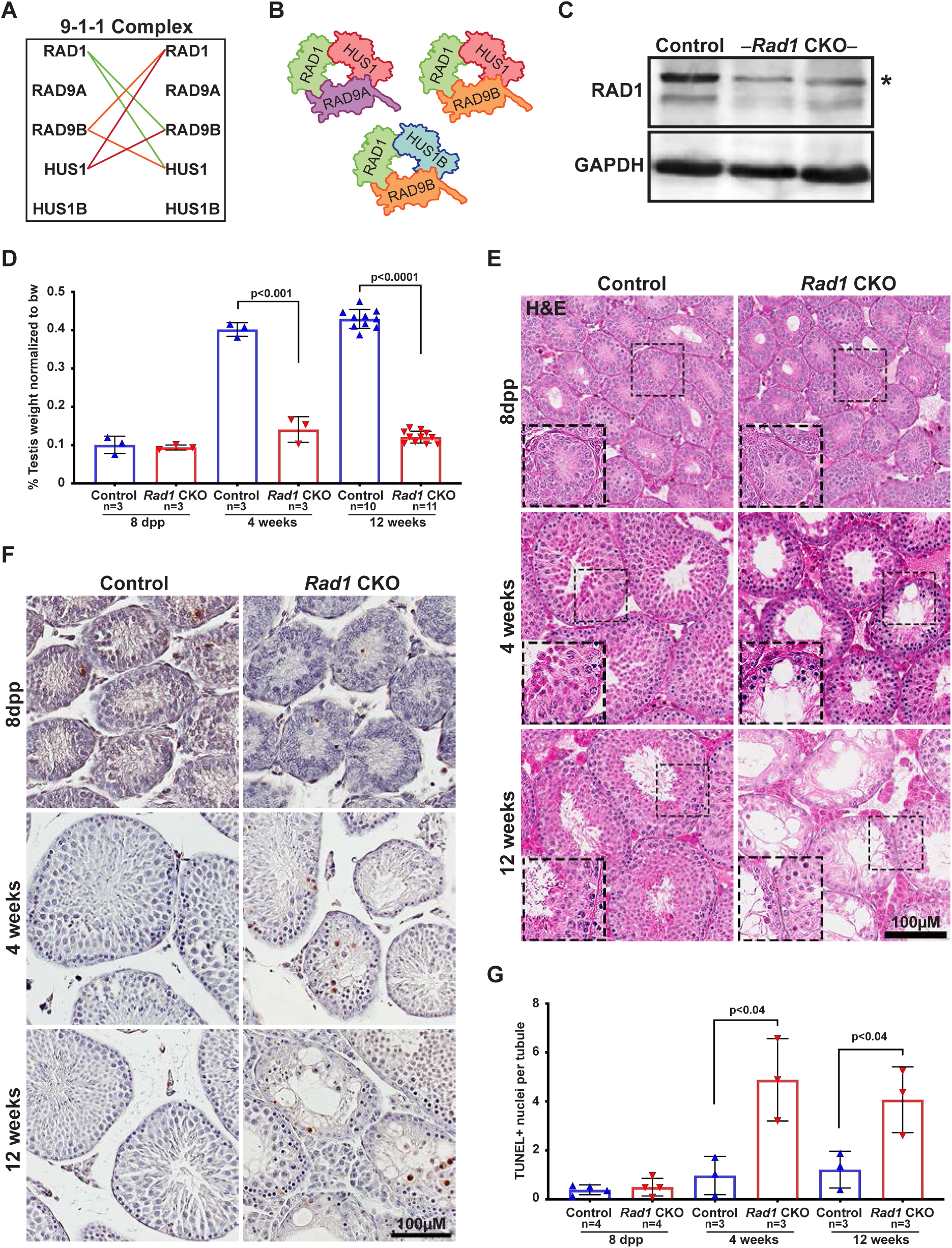
Conditional knockout targeting multiple 9-1-1 complexes causes severe germ cell loss in testes. (A) Evolutionary rate covariation analysis between 9-1-1 subunits. Lines depict significant covariance between 9-1-1 subunits. (B) Schematic showing putative meiotic 9-1-1 complexes: 9A-1-1, 9B-1-1 and 9B-1-1B. (C) Immunoblotting for RAD1 in control and *Rad1* CKO whole testes lysates from 12-week-old mice. (D) Testis weight normalized to body weight from 8-dpp, 4-week-old, and 12-week-old control and *Rad1* CKO mice. (E) Seminiferous tubule cross sections from 8 day postpartum (dpp), 4-week-old and 12-week-old mice were stained with H&E (representative images from 3 mice analyzed per age group per genotype). (F-G) Representative images (F) and quantification (G) of TUNEL positive cells per tubule in control and *Rad1* CKO mice (50 tubules per mouse quantified; n= number of mice analyzed). p-value calculated using Welch’s unpaired t-test using Graphpad.

### Testis-specific RAD1 loss leads to increased germ cell apoptosis and infertility

To determine how disrupting the subunit shared by all of the 9-1-1 complexes impacted meiosis, we created a *Rad1* CKO model by combining a conditional *Rad1* allele (Wit et al., 2011) with *Stra8-Cre*, which drives CRE expression in spermatogonia (Sadate-Ngatchou et al., 2008). A similar approach was previously used to create *Hus1* CKO mice (Lyndaker et al., 2013a), also on the inbred 129Sv/Ev background, enabling direct comparison of results between the two models. Experimental *Rad1* CKO mice carried one *Rad1^flox^* allele, one *Rad1-*null allele, and *Stra8-Cre* (*Rad1^-/fl^; Cre^+^*). Mice that carried a wild-type *Rad1* allele (*Rad1^+/fl^; Cre^+^*) or lacked *Stra8-Cre* (*Rad1^-/fl^; Cre^-^* or *Rad1^+/fl^; Cre^-^*) were used as littermate controls. Both *Rad1* CKO and control mice were born at expected frequency.

Immunoblotting of whole testis lysates from 12-week-old *Rad1* CKO mice confirmed significant reduction in RAD1 protein (n=5 control and 5 CKO; Figure 2C). The residual RAD1 protein observed in *Rad1* CKO mice could arise from somatic cells of the testis or pre-meiotic germ cells. However, we cannot exclude the possibility that persistent RAD1 protein exists in spermatocytes due to partial CRE recombinase efficacy or perdurance of RAD1 protein from pre-meiotic stages. Testes from *Rad1* CKO males were one-third the size of control testes at 4 weeks of age, while bodyweight was not altered (Figure 2D). Hematoxylin and eosin (H&E) staining of testis sections from control and *Rad1* CKO mice showed a reduction in tubule size starting at 4 weeks in CKO mice, with the phenotype being much more severe in 12-week-old mice (Figure 2D-E). Similar to previous findings in *Hus1* CKO males (Lyndaker et al., 2013a), *Rad1* CKO mice displayed increased apoptosis of zygotene/pachytene-staged cells (Figure 2F-G). In *Rad1* CKO mice, round spermatids were observed in some histology sections in 4-week-old and 12-week-old mice, possibly reflecting incomplete deletion and continued RAD1 expression in some meiocytes.

TUNEL staining confirmed significantly increased apoptosis in testes from *Rad1* CKO mice starting at 4 weeks of age (Figure 2F-G). 4-week-old *Rad1* CKO mice contained 2.4 ± 0.8 apoptotic nuclei per seminiferous tubule, compared to 0.5 ± 0.4 in control mice. Apoptosis continued to be significantly elevated in 12-week-old *Rad1* CKO mice (2.0 ± 0.7 positive nuclei per tubule) as compared to control mice (0.6 ± 0.4 positive nuclei per tubule) and was apparent in zygotene/pachytene-staged cells (Figure 2F-G). To quantify the impact of *Rad1* loss on germ cells, we stained testis sections for the germ cell-specific antigen TRA98 (Carmell et al., 2016). Tubules from control mice at 4 or 12 weeks of age contained an average of 220.2 ± 26.3 or 254.3 ± 45.5 TRA98-positive cells per tubule respectively (Figure 2-figure supplement 1A-B). However, in the absence of RAD1 tubules contained only 74.89 ± 7.5 TRA98-positive cells in 4-week-old mice and 47.8 ± 8.28 in 12-week-old males.

*Stra8*-*Cre* expression occurs as cells are committing to undergo meiosis (Sadate-Ngatchou et al., 2008). We therefore anticipated that the apoptosis and germ cell loss observed in *Rad1* CKO mice were due to meiotic defects. To address the possibility of pre-meiotic defects in *Rad1* CKO mice, we assessed mice at 8 days postpartum (dpp), prior to meiotic entry. H&E staining, along with TUNEL and TRA98 staining of sections from both control and *Rad1* CKO mice, showed no significant differences between genotypes at 8 dpp (Figure 2-figure supplement 1A-D). To further confirm that RAD1 loss did not affect cells prior to meiotic entry, we stained sections for LIN28, a marker of spermatogonial stem cells (SSCs), which have not initiated meiosis (Aeckerle et al., 2012). As expected, no significant differences in LIN28 staining were observed between genotypes in testes from mice at 8 dpp or 4 weeks of age (Figure 2-figure supplement 1C-D), consistent with the notion that RAD1 targeting is specific to meiotic cells. However, 12-week-old *Rad1* CKO mice had a significant decrease in LIN28-positive cells when compared to control mice. This later loss of LIN28-positive cells in *Rad1* CKO mice can be attributed to the large-scale germ cell loss, which could indirectly disrupt the environment required for proper SSC proliferation and survival.

Next, we tested how localization of 9-1-1 subunits was affected by RAD1 loss. Consistent with prior results (Freire et al., 1998; Lyndaker et al., 2013a) RAD1 localized in control mice as foci on chromosome cores that were not yet synapsed during leptonema and zygonema, and was present on the core axis of the X and Y chromosomes in pachynema (Figure 3A). RAD1 expression was completely absent in 43% of spermatocytes from 12-week-old *Rad1* CKO mice, whereas 100% of control cells showed proper RAD1 localization and abundance in zygotene-and pachytene-stage cells (Figure 3A). RAD1 expression in *Rad1* CKO meiocytes could be attributed to cells that failed to undergo CRE-mediated recombination or in which RAD1 levels were not yet fully depleted. The fact that *Rad1* CKO cells were prone to apoptosis would be expected to eliminate cells lacking RAD1, leaving RAD1-intact meiocytes enriched among the remaining cells. We next evaluated how RAD1 loss impacted RAD9A/B localization. In control samples, RAD9A and RAD9B localized to unsynapsed regions as foci in leptotene-and zygotene-staged cells (Figure 3B and Figure 3-figure supplement 1A). By pachynema, RAD9A and RAD9B localized primarily to the XY cores as foci. RAD9A and RAD9B localization were absent in 79% and 72%, respectively, of *Rad1* CKO meiotic spreads (Figure 3B and Figure 3-figure supplement 1A).

**Figure 3:**
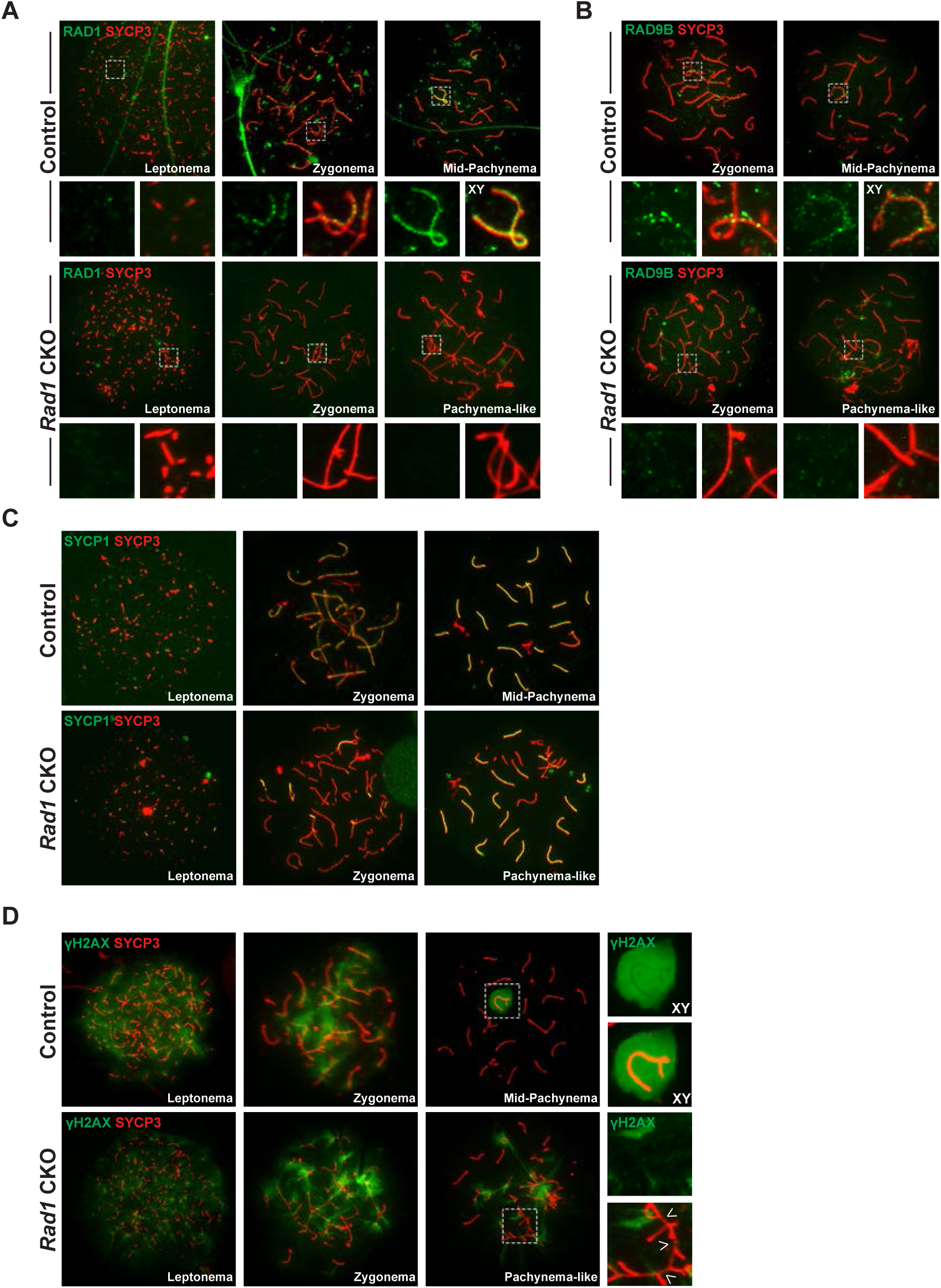
Increased asynapsis and DNA damage is observed upon testis-specific RAD1 loss. (A-B) Meiotic spreads from 12-week-old control and *Rad1* CKO mice stained for RAD1 (A) or RAD9B (B). (C) Co-staining of SYCP1 and SYCP3 from 12-week-old control and *Rad1* CKO mice (3 control mice, n= 156 cells; 3 CKO mice, n= 131 cells). *Rad1* CKO meiocytes with 4 or more synapsed chromosomes were categorized as pachytene-like. (D) **γ**H2AX staining of meiotic spreads from control and *Rad1* CKO mice. Arrowheads in *Rad1* CKO spreads highlight regions of asynapsis lacking **γ**H2AX staining (3 control mice, n= 127 cells; 5 CKO mice, n= 205 cells). P-values were calculated using Welch’s unpaired t-test using Graphpad.

*Rad1* CKO mice had no epididymal sperm (Table 1). To assess if *Rad1* CKO mice were infertile, control and *Rad1* CKO mice were bred with wild-type females. Control mice bred with wild-type females yielded 10 pregnancies and 66 viable pups, whereas *Rad1* CKO mice had no viable offspring from 15 matings with wild-type females. Overall, these results indicate that RAD1 disruption severely compromised spermatogenesis and fertility. Moreover, the reduced testis weight and increased apoptosis in *Rad1* CKO mice were more severe than those in mice with *Hus1* or *Rad9a* loss (Lyndaker et al., 2013a; Vasileva et al., 2013), suggesting a broader role for RAD1 in meiocytes.

**Table 1:** Analysis of epididymal sperm counts and fertility in *Rad1* CKO and control mice^1^.

| Genotype | # males | Epididymal Sperm Count (x10<sup>6</sup>) | # matings | # copulatory plugs | # pregnancies | Total viable pups |
| --- | --- | --- | --- | --- | --- | --- |
| Control | 3 | 16.6 ± 4.5 | 12 | 12 | 10 | 66 |
| *Rad1* CKO | 3 | 0.0 ± 0 | 15 | 15 | 0 | 0 |

<sup>1</sup> Male *Rad1* CKO mice at 8 to 12 weeks of age were bred to 6-week-old wild-type FVB female mice.

### *Rad1* loss results in synapsis defects and increased DNA damage

During meiosis, SC formation is critical for homologous chromosomes to pair and fully synapse (Zickler and Kleckner, 2015). Co-staining for the SC markers SYCP1 and SYCP3 revealed that 59.5 ± 4.3% of meiocytes from *Rad1* CKO mice had whole chromosomes that remained unsynapsed and/or aberrant synapsis events involving multiple chromosomes, whereas 100% of meiocytes from control mice displayed normal homolog synapsis (Figure 3C and Figure 3-figure supplement 1C). RAD1 staining in meiocytes from 12-week-old *Rad1* CKO mice revealed that all cells that lacked RAD1 displayed abnormal synapsis, with an average of only 8 chromosomes fully synapsed chromosomes per cell (Figure 3-figure supplement 1D). Cells with asynapsis that contained four or more synapsed homologous chromosomes were classified as pachytene-like cells, and subsequent analyses focused on this population of *Rad1* CKO meiocytes.

The γH2AX staining pattern was similar in *Rad1* CKO and control spreads at leptonema and zygonema (Figure 3D). However, 97% of pachytene-like *Rad1* CKO cells showed γH2AX present at asynaptic sites, with no clear presence of a sex body (n= 98 cells, 3 CKO mice). Interestingly, a subset of asynaptic regions in *Rad1* CKO cells lacked detectable γH2AX staining (Figure 3D, white arrow heads), suggesting that the DNA damage signaling at asynaptic sites was perturbed (Figure 3-figure supplement 1B).

Given that spermatocytes from *Rad1* CKO mice exhibited significantly increased asynapsis, we next assessed meiotic progression in these cells by staining for the histone variant H1T and the recombination marker MLH1. First, we questioned whether *Rad1* CKO cells were able to progress past mid-pachynema. Histone variant H1T is a marker of mid-pachynema and later staged wild-type spermatocytes (Barchi et al., 2005). Control cells demonstrate H1T staining as they progress into mid-pachynema (Figure 3-figure supplement 1E). However, H1T staining was absent in *Rad1* CKO meiocytes, indicating that the cells failed to progress past mid-pachynema (Figure 3-figure supplement 1E). By mid-pachynema, crossover sites are normally marked by MLH1 (Eaker et al., 2002). MLH1 was not detected in any *Rad1* CKO cells at the pachytene-like stage (Figure 3-figure supplement 1F), further suggesting that *Rad1* CKO meiocytes fail to progress beyond early/mid-pachynema. Together the observations of γH2AX abnormalities and SC defects in *Rad1* CKO cells indicate important roles for 9-1-1 complexes in ensuring homologous chromosome synapsis and appropriate DDR signaling in response to asynapsis.

### DSB repair is compromised in *Rad1* CKO spermatocytes

Because testis-specific *Hus1* or *Rad9a* CKO results in persistent meiotic DSBs with delayed repair kinetics (Lyndaker et al., 2013a; Vasileva et al., 2013), we investigated how RAD1 loss impacts DSB repair. Following MRE11-RAD50-NBS1 (MRN)-mediated resection of SPO11-induced meiotic DSB, Meiosis-specific with OB domains (MEIOB) and RPA localize to the ssDNA overhangs prior to RAD51 and DMC1 loading (Hinch et al., 2020; Luo et al., 2013; Shi et al., 2019). In control spermatocytes, RPA and MEIOB foci are abundant in early prophase I and diminish as DSBs are repaired. *Rad1* CKO testes had on average 50 fewer RPA foci than controls in leptotene-stage cells (194 ± 54 control; 146 ± 37 CKO; Figure 4A-B). Intriguingly, RPA foci in *Rad1* CKO cells appeared larger than those in control cells. In the absence of RAD1, MEIOB focus formation on chromatin cores in leptotene-stage cells was also significantly decreased as compared to control cells (230 ± 45 control; 126 ± 37 CKO; Figure 4C-D). In control spermatocytes, MEIOB and RPA levels on meiotic chromosome cores decreased as the cells progressed into pachynema (115 ± 27 control MEIOB; 52 ± 41 control RPA), whereas *Rad1* CKO cells showed persistence of MEIOB and RPA staining (132 ± 38 CKO MEIOB; 100 ± 44 CKO RPA; Figure 4A-D).

**Figure 4:**
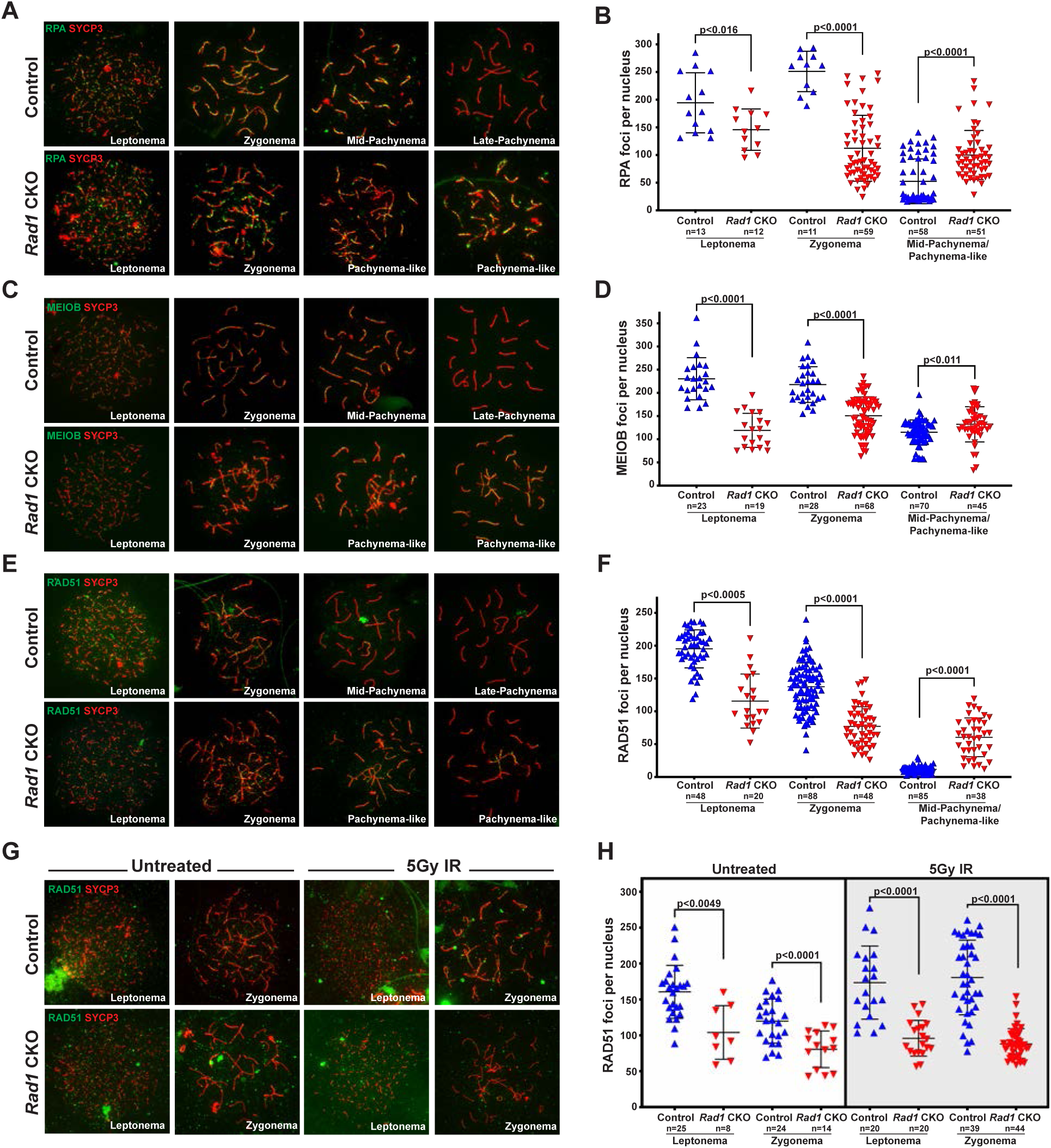
DSB repair is compromised in the absence of 9-1-1 complexes. (A-B) Representative images (A) and quantification (B) of RPA staining of meiotic spreads from 12-week-old control and *Rad1* CKO mice (3 mice per genotype analyzed; n=total cells analyzed). (C-D) Representative meiotic spread images for ssDNA marker MEIOB (C) and quantifications (D) from 12-week-old control and *Rad1* CKO mice (3 mice per genotype analyzed; n= total cells analyzed). (E-F) Representative meiotic spread images of RAD51 (E) and quantifications (F) from 12-week-old control and *Rad1* CKO mice (5 control mice and 6 CKO analyzed; n=total cells analyzed). (G-H) 8-week-old control and *Rad1* CKO mice were irradiated with 5Gy IR and collected 1-hour post IR. Representative RAD51 meiotic spread images (G) and quantifications (H) (2 control mice and 2 CKO analyzed; n= total cells analyzed). P-values were calculated using Welch’s unpaired t-test using Graphpad.

During prophase I in wild-type spermatocytes, RAD51 and DMC1 displace MEIOB and RPA from the ssDNA overhangs and drive the subsequent steps of homology search and strand invasion (Hinch et al., 2020; Luo et al., 2013; Shi et al., 2019). The persistence of MEIOB and RPA foci in *Rad1* CKO spermatocytes suggested that RAD1 loss might perturb RAD51 loading. On average, leptotene-stage cells from control mice contained 195 ± 29 RAD51 foci, whereas *Rad1* CKO cells at the same stage had 60 ± 30 RAD51 foci (Figure 4E-F). RAD51 foci continued to be significantly lower in zygotene-stage *Rad1* CKO meiocytes, which contained 96 ± 43 RAD51 foci per cells as compared to 146 ± 31 in controls. In control samples, RAD51 foci levels decreased as cells progressed from zygonema to pachynema, reflecting the successful repair of DSBs. However, *Rad1* CKO spermatocytes retained relatively high levels of RAD51 foci in pachytene-like-stage cells (67 ± 44 RAD51 foci) as compared to control pachytene-stage meiocytes (10 ± 4 RAD51 foci) (Figure 4F). These results for RAD51 localization in *Rad1* CKO spermatocytes differed from those in *Hus1* CKO mice, where RAD51 appeared normal in early prophase and then was aberrantly retained at a small number of sites in pachytene-stage cells (Lyndaker et al., 2013a). Together these results suggest that the 9-1-1 complexes are critical for proper DSB repair during mammalian meiosis and that absence of RAD1, or to a lesser extent HUS1, leaves persistent unrepaired DSBs.

The delayed loading of MEIOB, RPA and RAD51 observed in *Rad1*-deficient spermatocytes raised the possibility that DSB formation was impaired. To determine whether the defects were related to DSB formation or the subsequent repair steps, we treated *Rad1* CKO and control mice with 5 Gy ionizing radiation, harvested testes 1-hour post treatment, and quantified RPA and RAD51 focus formation in leptotene-and zygotene-stage cells. Since exogenously induced DSBs are repaired via meiotic processes in early stages of prophase I (Enguita-Marruedo et al., 2019), this approach allowed us to test whether the alterations in DSB markers in *Rad1* CKO cells were due to reduced DSB formation or a DSB repair defect. Upon DSB induction via irradiation, control mice showed the expected increase in RPA and RAD51 focus formation at early prophase I stages (Figure 4G-H, and Figure 4-figure supplement 1A-B). By contrast, irradiation did not induce increased focus formation by RPA or RAD51 in *Rad1* CKO spermatocytes. These results suggest an intrinsic defect in meiotic DSB repair when the 9-1-1 complexes are disabled.

### The localization of key components of the ATR axis is compromised in the absence of RAD1

ATR is a primary regulator of MSCI, which involves several DDR factors that are ATR substrates (Pacheco et al., 2018; Turner, 2007, 2015; Widger et al., 2018). Given that the canonical 9A-1-1 complex plays a central role in stimulating ATR activity in somatic cells, we sought to determine the effect of RAD1 loss on the localization of ATR and its substrates in meiocytes. ATR localizes to unsynapsed regions at early stages of prophase I, and by pachynema it is sequestered mainly at the XY body where it initiates MSCI (Abe et al., 2020; Turner, 2015). Cells from *Rad1* CKO mice with synapsis defects showed ATR localization only at a subset of unsynapsed regions (Figure 5A).

**Figure 5:**
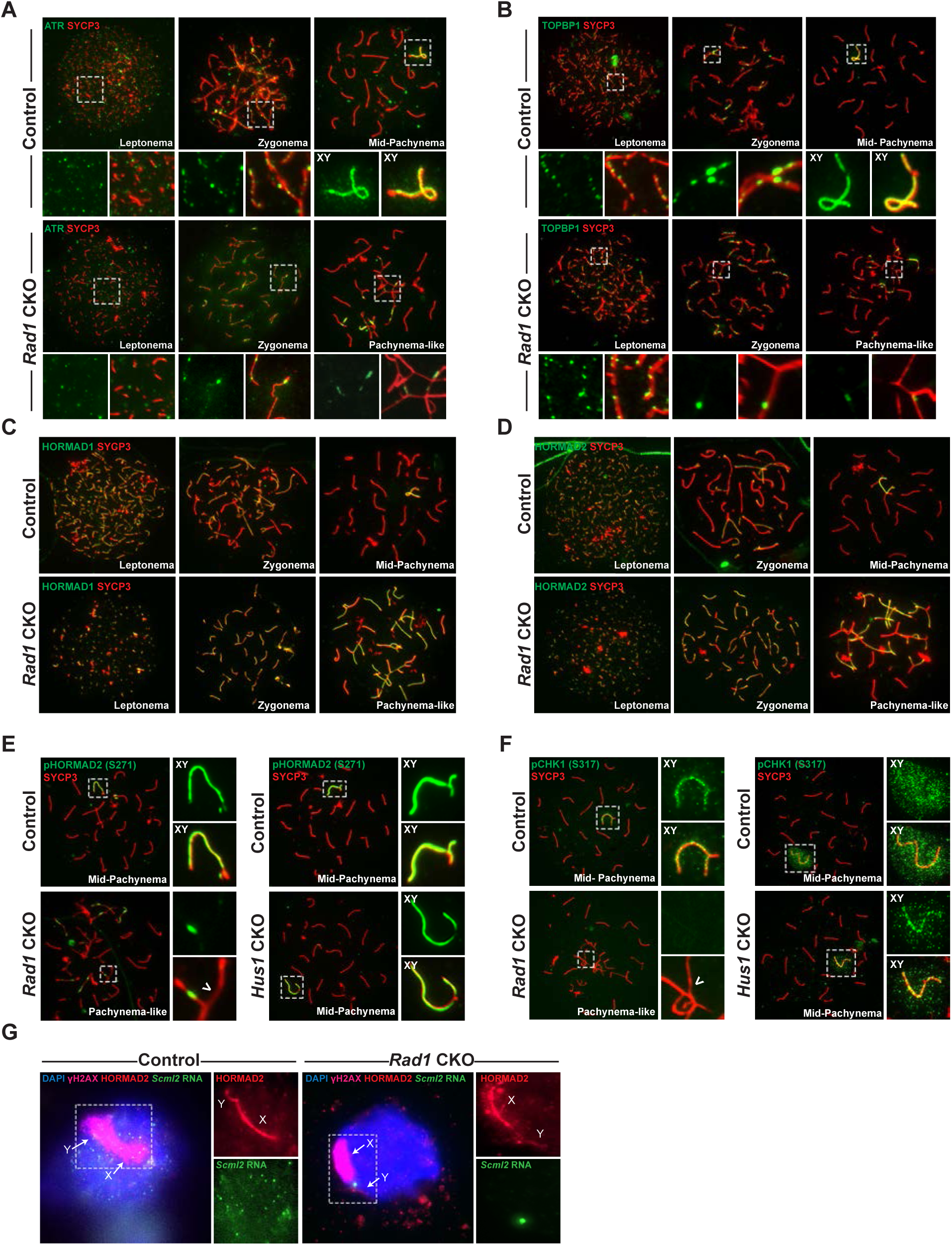
ATR localization and function is dependent upon 9-1-1 complexes. (A) ATR localization in control and *Rad1* CKO 12-week-old meiotic spreads (3 control, n= 171 cells; 3 CKO, n= 146). (B) Representative images of TOPBP1 localization in meiotic spreads from 12-week-old control and *Rad1* CKO mice (3 control, n= 130 cells; 3 CKO, n= 129). (C-D) Representative images of HORMAD1 and HORMAD2 localization in 12-week-old control and *Rad1* CKO mice (3 control, n= 146; 3 CKO, n= 119). (E) Representative images of phospho-HORMAD2 (S271) localization in 12-week-old control and *Rad1* CKO mice (*Rad1* CKO: 3 control, n= 178; 3 CKO, n= 146. *Hus1* CKO: 2 control, n= 189; 3 CKO, n= 145). (F) Representative images of phospho-CHK1 (S317) localization in *Rad1* CKO and *Hus1* CKO mice (*Rad1* CKO: 3 control, n= 125; 3 CKO, n= 120. *Hus1* CKO: 2 control, n= 107; 3 CKO, n= 191). (G) RNA FISH for *Scml2* in fully synapsed, pachytene-stage control and *Rad1* CKO cells (3 control, n= 29 cells; 3 CKO, n= 45 cells).

TOPBP1 is required for ATR activation following replication stress (Mordes et al., 2008) and interacts with ATR during meiosis to ensure that meiotic silencing is properly initiated (ElInati et al., 2017; Jeon et al., 2019). In control meiocytes, TOPBP1 was observed as discrete foci on unsynapsed chromosome cores throughout leptonema and zygonema (Figure 5B). At pachynema, TOPBP1 was found exclusively along the unsynapsed regions of the X and Y and present as a faint cloud on XY chromosome loops. By contrast, in pachytene-like stage *Rad1* CKO cells, TOPBP1 localized to only a subset of asynaptic sites, failing to coat the entirety of unsynapsed chromosome cores, similar to the pattern observed for ATR. Although these findings may suggest a role for the 9-1-1 complexes in promoting ATR and TOPBP1 localization to unsynapsed chromosomal regions, it remains possible that the extensive asynapsis in *Rad1* CKO mice causes an insufficiency in the available pool of silencing factors needed to localize to all asynaptic sites (Mahadevaiah et al., 2008).

Localization of HORMA-domain proteins 1 and 2 (HORMAD1 and HORMAD2) at unsynapsed chromatin is important for meiotic silencing (Fukuda et al., 2010; Wojtasz et al., 2009). Furthermore, HORMAD1 plays an important role in ATR recruitment to unsynapsed sites (Fukuda et al., 2010; Shin et al., 2010). We next examined whether defective ATR localization in the absence of RAD1 was the result of HORMAD mislocalization. In control cells, HORMAD1 and HORMAD2 were observed during early prophase I at chromosomal regions that were not yet synapsed (Figure 5C-D). By mid-pachynema the HORMADs localized strictly at the unsynapsed regions of the XY, similar to the localization of ATR. Notably, RAD1 loss did not alter HORMAD1 or HORMAD2 localization to unsynapsed regions. Furthermore, the HORMADs were observed to entirely coat unsynapsed chromosome regions in *Rad1* CKO cells, in contrast to the failure of ATR and TOPBP1 to localize to all unsynapsed sites.

### Phosphorylation of key ATR substrates is disrupted in Rad1 CKO meiocytes

ATR phosphorylates HORMAD1 (S375) and HORMAD2 (S271) at asynaptic regions (Fukuda et al., 2012; Royo et al., 2013). In control cells, HORMAD2 (S271) phosphorylation was observed on the X and Y chromosome cores in mid-pachytene-stage cells as expected (Figure 5E). However, in pachytene-like *Rad1* CKO cells, phosphorylated HORMAD2 was detected at only a subset of unsynapsed regions. That HORMAD2 localized properly in the absence of RAD1 but lacked phosphorylation at an ATR-regulated site strongly suggests a requirement for 9-1-1 complexes in meiotic ATR signaling.

The best characterized ATR substrate in somatic cells is the transducer kinase CHK1. CHK1 has been proposed to play a role in meiotic DSB repair and is suggested to aide progression through prophase I by removal of DNA damage response proteins such as **γ**H2AX from autosomes {Abe, 2018 #163;Fedoriw, 2015 #157;Pacheco, 2018 #126}. In wild-type cells, CHK1 phosphorylation (S317) occurs during leptonema and zygonema at unsynapsed chromosomes. During pachynema, pCHK1 (S317) is apparent as a cloud over the sex body, similar to **γ**H2AX and ATR (Figure 5F). Interestingly, in the *Rad1* CKO mutant, pCHK1 was absent at all stages of prophase I. By contrast, meiotic spreads from *Hus1* CKO mice showed normal patterns of CHK1 (S317) and HORMAD2 (S271) phosphorylation (Figure 5E-F). That meiotic CHK1 phosphorylation is normal in the absence of HUS1 but disrupted by RAD1 loss suggests that alternative 9-1-1 complexes play an important role in activating the transducer kinase CHK1 during meiotic prophase I.

The defects in ATR signaling observed in *Rad1* CKO mice suggested that disruption of 9-1-1 complexes might impair meiotic silencing. To test this possibility, we evaluated meiotic silencing via RNA fluorescent in situ hybridization (FISH) for the X-chromosome gene *Scml2* that should be silenced in early pachynema-stage cells (Royo et al., 2010). *Scml2* expression was detected in 7.1 ± 0.6% of early pachytene control cells while *Rad1* CKO cells showed expression in 28.9 ± 3.2% (p<0.0001; Figure 5G), indicating that meiotic silencing was disrupted by RAD1 loss. The analysis of *Scml2* expression focused on cells with normal homolog synapsis and excluded those with asynapsis, which could underestimate the extent of the silencing defect upon RAD1 loss since some cells with normal synapsis in *Rad1* CKO mice retain RAD1 expression. Nevertheless, these data demonstrate the importance of alternative 9-1-1 complexes in ensuring that ATR-mediated MSCI occurs.

### Comprehensive profiling of protein phosphorylation in testes from *Rad1* CKO mice

Our findings that phosphorylation of key ATR substrates like HORMAD2 and CHK1 was disrupted in *Rad1* CKO mice prompted us to more thoroughly characterize how RAD1 loss impacts meiotic signal transduction. Since the 9-1-1 complex is well-established to regulate ATR activation, we sought to identify phosphorylation events that were dependent on both the 9-1-1 complex and ATR. To accomplish this, we analyzed not only *Rad1* CKO testes but also those from wild-type C57BL/6 (B6) mice treated with the ATR inhibitor (ATRi) AZ20 (Sims et al., 2021). Wild-type B6 males received either chronic (3 doses of 50mg/kg over 3 days) or acute (single dose of 50mg/kg for 4 hrs) ATR inhibitor treatment. Whole testis lysates were subjected to phosphoproteomic analysis by mass spectrometry. The *Rad1* CKO samples featured cell type-specific disruption of RAD1, but the chronic nature of RAD1 depletion might lead to secondary alterations in the phosphoproteome of these cells. Though not cell-type specific, systemic ATR inhibition was acute, limiting secondary effects. The combined phosphoproteomic analysis of *Rad1* CKO and ATRi-treated testes allowed us to accurately identify meiotic phosphorylation events dependent on the combined actions of the 9-1-1 complexes and ATR (Figure 6A-B).

**Figure 6:**
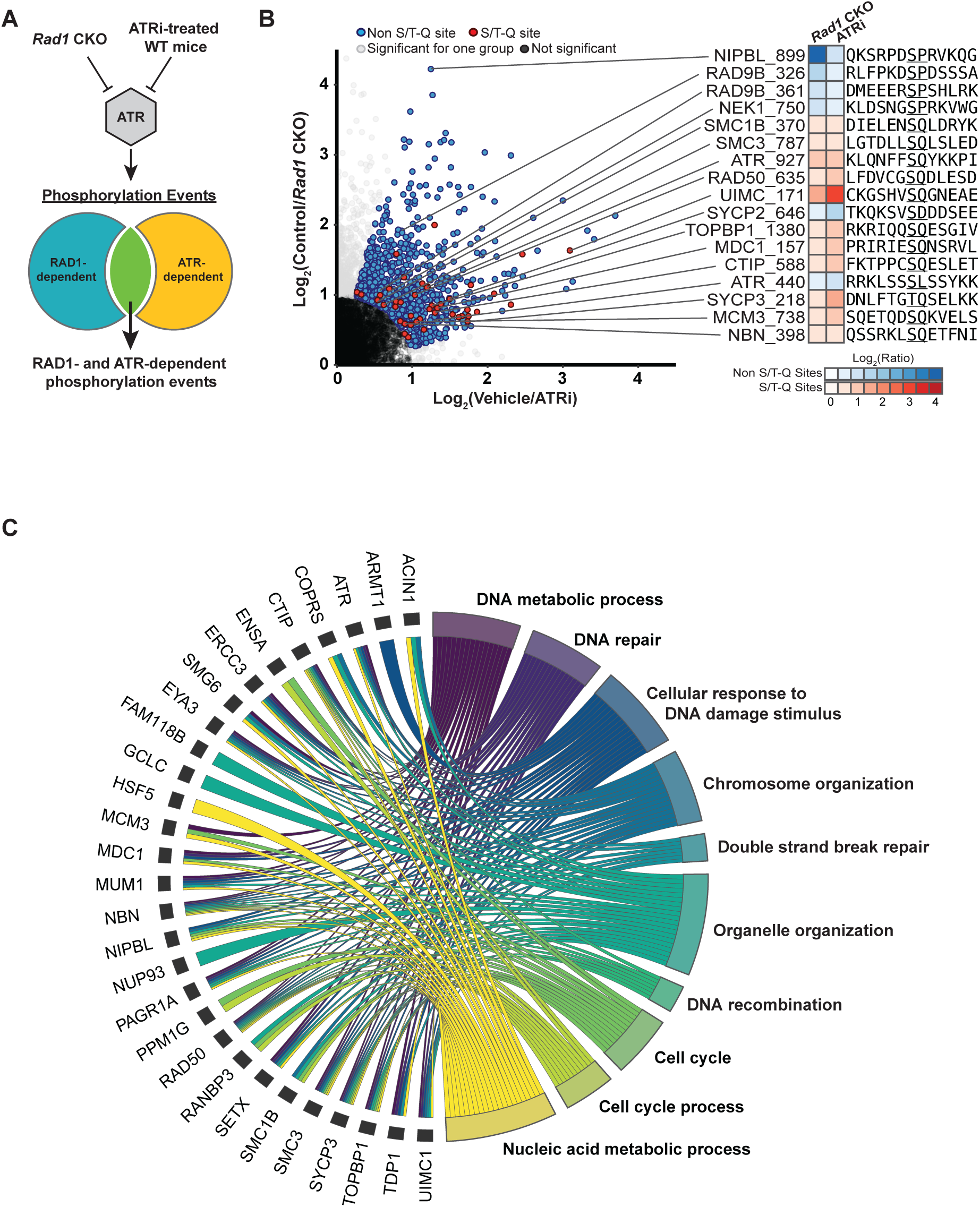
Coordinated roles for 9-1-1 complexes and ATR in phospho-regulation of DDR and cohesin proteins. (A) Decapsulated testes from adult *Rad1* CKO mice or WT C57BL/6 mice treated with ATRi were subjected to phosphoproteome analysis. Subsequent analyses focused on proteins phosphorylated in a RAD1-and ATR-dependent manner. (B) Bowtie analysis showed a total of 863 ATR/RAD1-dependent phosphorylation events detected. Red circles indicate S/T-Q sites (42 peptides), blue are non-S/T-Q sites, gray are significant phosphosites in either only the ATRi or the *Rad1* CKO experiment, and black circles were phosophsites in both experiments but were not differentially phosphorylated. For the indicated phosphoproteins, the heatmap shows the average of the log2 fold change across replicates for each experiment (ATRi and *Rad1* CKO), and the amino acid sequence shows 6 residues upstream and 6 residues downstream from the indicated phosphosite. (C) Gene ontology analysis of ATR/RAD1-dependent S/T-Q phosphorylation events was done using STRINGdb. The top ten significantly enriched biological processes GO terms were selected and represented as a chord diagram. GO terms are shown on the right and proteins found for each term on the left.

Principal component analysis showed tight clustering of three independent *Rad1* CKO samples and established that the *Rad1* CKO samples differed from ATRi samples along principal components 1 and 2 (Sims et al., 2021). High-quality phosphopeptides were designated after filtering for sites with a localization score >0.85 and considering only sites found in at least two independent experiment. Filtering of the data in this manner resulted in a total of 12,220 phosphopeptides. 863 phosphopeptides had significantly reduced phosphorylation in both *Rad1* CKO and ATRi samples relative to their matched controls and were considered to be RAD1-and ATR-dependent. 42 of these were differentially phosphorylated at S/T-Q sites, the ATR target motif (Blackford and Jackson, 2017) (Figure 6B).

STRING-db was used to obtain gene ontologies (GO) for the differentially phosphorylated S/T-Q sites, and the top ten terms were highlighted in a chord diagram (Figure 6C). Major pathways known to be linked to ATR and the 9-1-1 complex were identified, with the top two terms being cellular response to DNA metabolic processes and DNA repair. The GO chord diagram further showed that nine of the terms included TOPBP1, and six of the terms included ATR. DDR proteins whose phosphorylation was dependent upon both RAD1 and ATR included proteins such as MDC1, involved in meiotic silencing, as well as DNA end-resection factors like CTIP and RAD50. The reduced phosphorylation of CTIP and RAD50 was notable in light of the observation that localization of the ssDNA binding proteins RPA and MEIOB, as well as the recombinase RAD51, was defective in *Rad1* CKO meiocytes, hinting at a potential role for the 9-1-1 complexes in DSB processing.

### A role for 9-1-1 complexes in cohesin regulation

Cohesins are critical for ensuring proper chromosome segregation in both mitotic and meiotic cells (Ishiguro, 2019). Loss of meiosis-specific cohesins, such as SMC1β, REC8 or RAD21L, results in phenotypes that include DSB repair failure and synapsis defects (Challa et al., 2019; Ishiguro, 2019; Ward et al., 2016). SC assembly is also dependent upon proper cohesin loading (Eijpe et al., 2003; Llano et al., 2012). Interestingly, phosphorylation of SC components SYCP1 and SYCP2 was reduced in both *Rad1* CKO and ATRi-treated mice in the phosphoproteomic screen (Figure 6B-C). Cohesin complex components such as SMC3 and SMC1β also showed reduced phosphorylation in both ATRi and *Rad1* CKO samples. Furthermore, correlated evolutionary relationships, as measured by ERC analysis, were observed between genes encoding 9-1-1 subunits and those encoding cohesin and SC proteins, including SMC1β, RAD21L1, SYCP2, and SYCE1 (Figure 7A-B and Figure 7-figure supplement 1A-B). ERC network analysis of the relationship between proteins involved in meiosis I and the 9-1-1 subunits revealed a clustering of RAD9B, RAD1 and HUS1, while RAD9A and HUS1B did not show high ERC values with the other 9-1-1 subunits and had mostly separate network interactions (Figure 7-figure supplement 1A-B). Both the ERC data and phosphoproteomic results implicate RAD1-containing 9-1-1 complexes in SC formation and cohesin during mammalian meiosis, consistent with the aberrant synapsis observed in *Rad1* CKO but not *Hus1* CKO spermatocytes.

**Figure 7:**
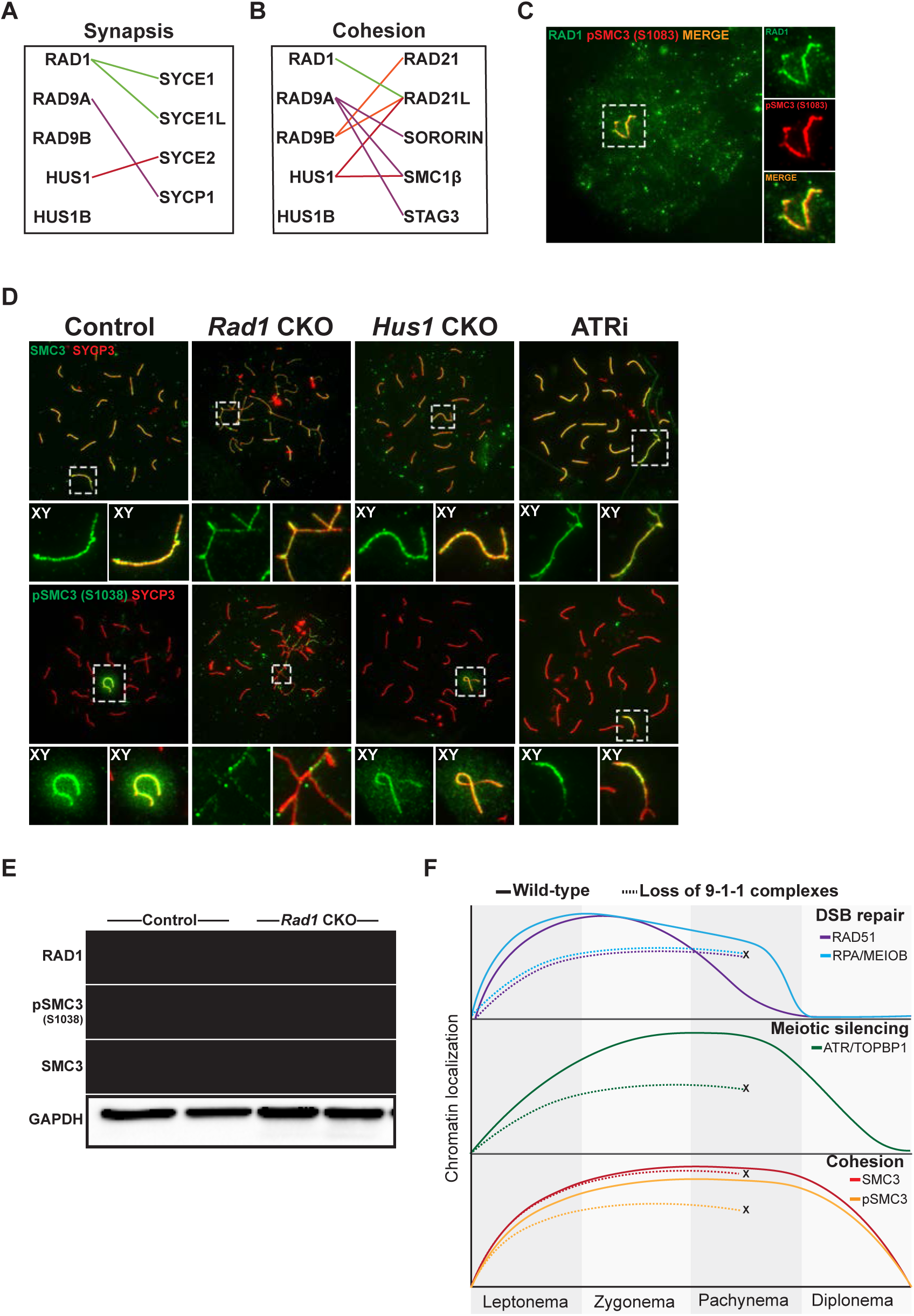
Cohesin phosphorylation events are dependent upon the 9-1-1/ATR axis. (A-B) ERC analysis between 9-1-1 complex subunits and synaptonemal complex (A) and cohesin (B) factors. Lines depict significant correlations observed between 9-1-1 complex subunits and synapsis or cohesin factors. (C) Co-staining of RAD1 and pSMC3 (1083) in wild-type spermatocytes. (D) Representative images of SMC3 and pSMC3 (1038) localization in pachynema and pachytene-like cells from control, *Rad1* CKO, *Hus1* CKO and ATRi-treated mice. (E) Representative immunoblot for phosphorylated SMC3 and total SMC3 in whole testes lysates from 12-week-old control and *Rad1* mice. (F) Summary graphic depicting the localization of key meiotic factors in wild-type versus *Rad1* CKO spermatocytes. Loss of 9-1-1 complexes resulted in early-to-mid pachynema arrest, as depicted by the ‘X’. DSB repair markers, such as RAD51, were reduced in the absence of the 9-1-1 complexes. Meiotic silencing factors such as ATR and TOPBP1 also failed to localize properly in the absence of the 9-1-1 complexes. The cohesin subunit SMC3 localized properly in in the absence of 9-1-1 subunits but its phosphorylation was impaired in *Rad1* CKO spermatocytes.

SMC3 phosphorylation (pSMC3) was previously shown to be ATR-dependent throughout meiotic prophase (Fukuda et al., 2012). Co-staining for RAD1 and pSMC3 (1083) in wild-type spermatocytes revealed co-localization of RAD1 and pSMC3 (1083) at the XY in pachynema-stage cells (Figure 7C). In control meiocytes, SMC3 was observed on chromatin cores throughout prophase I and was phosphorylated specifically at unsynapsed chromatin cores during leptonema and zygonema, and at the unsynapsed regions of the XY in mid-pachynema (Figure 7D). Although total SMC3 loading was unaffected, *Rad1* CKO spermatocytes showed reduced accumulation of phosphorylated SMC3 (pSMC3 S1083) at unsynapsed chromatin regions in pachytene-like cells as compared to mid-pachytene-stage control cells. Moreover, western blot analysis of whole testis lysates confirmed that SMC3 phosphorylation (pSMC3 S1083) was significantly reduced in testes from *Rad1* CKO mice (Figure7E). Unlike *Rad1* CKO spermatocytes, *Hus1* CKO cells had grossly normal pSMC3 (S1083) localization to the XY in pachytene-stage spermatocytes. Interestingly, chronic ATRi treatment caused a decrease in pSMC3 (S1083) localization to X and Y chromatin loops and XY cores (Figure 7D). Similar to the results in *Rad1* CKO spermatocytes, pSMC3 (S1083) localization was perturbed in spermatocytes from ATRi-treated mice despite the fact that SMC3 localization to chromosome cores appeared normal, suggesting a specific defect in SMC3 phosphorylation (Figure 7D top panel). Together these results suggest that 9-1-1 complexes and ATR act in conjunction to regulate meiotic cohesin phosphorylation.

## DISCUSSION

Here we report that testis-specific RAD1 loss results in defective homolog asynapsis, compromised DSB repair, faulty ATR signaling, and impaired meiotic silencing. Previous analyses of the canonical 9A-1-1 complex in meiosis revealed that loss of *Hus1* or *Rad9a* leads to a small number of unrepaired DSBs that trigger germ cell death (Lyndaker et al., 2013a; Vasileva et al., 2013). Yet, homolog synapsis, ATR activation and meiotic silencing all are grossly normal in the absence of the canonical 9-1-1 subunits HUS1 and RAD9A. The expanded roles for RAD1 identified here are consistent with its ability to additionally interact with RAD9B and HUS1B, paralogs that evolved in higher organisms and are highly expressed in germ cells. Together, our results support the idea that alternative 9-1-1 complexes evolved to play essential roles in meiotic DSB repair, homolog synapsis, and MSCI.

In *Rad1* CKO spermatocytes, RAD51 loading onto meiotic chromosome cores was significantly reduced at leptonema and zygonema relative to controls. In control cells, DSB repair is concluding and RAD51 chromatin levels are low by mid-pachynema, but substantial RAD51 focus formation was still observed in pachytene-like *Rad1* CKO cells, suggesting major DSB repair defects. The meiotic DSB repair defects following RAD1 loss are similar to those previously observed in *Atr* loss of function mouse models. Zygotene-stage cells from a Seckel mouse model with disrupted ATR expression have decreased RAD51 and DMC1 loading (Pacheco et al., 2018), similar to that of spermatocytes lacking RAD1. Meiotic RAD51 focus formation did not increase further in *Rad1* CKO meiocytes after irradiation. These finding suggest that, similar to what is observed in *Atr*-defective spermatocytes (Pacheco et al., 2018; Widger et al., 2018), the defects in RAD51 loading were not due to decreased numbers of SPO11-induced DSBs in *Rad1* CKO mice, highlighting an important role for the 9-1-1 complexes in the subsequent repair of meiotic DSBs.

Unlike what is observed in *Atr* mutants and ATR inhibitor-treated mice, localization of ssDNA markers MEIOB and RPA to meiotic cores was significantly reduced in the absence of RAD1. The 9-1-1 complex is well-established to modulate DNA end resection, having stimulatory or inhibitory effects in different contexts. In both yeast and mammals, the resection-stimulatory effects of the 9-1-1 complex involve recruitment of the Exo1 and Dna2 nucleases to DNA (Blaikley et al., 2014; Karras et al., 2013; Ngo et al., 2014; Ngo and Lydall, 2015). Our phosphoproteomic analysis of *Rad1* CKO testes and ATRi-treated mice also revealed a significant decrease in phosphorylation of proteins involved in DNA end resection, including RAD50, NBS1, and CTIP. Conditional *Nbs1* knockout in testes was previously reported to cause a decrease in chromatin loading of RPA, MEIOB and RAD51 (Zhang et al., 2020), similar to that in *Rad1* CKO mice, further suggesting potential functional interplay between the 9-1-1/ATR signaling axis and MRN complex during meiosis.

Somatic activation of ATR via 9A-1-1/TOPBP1 interaction is well established; however, ATR and TOPBP1 localization in spermatocytes was unperturbed in the absence of *Hus1*. ATR-dependent processes such as sex body formation and meiotic silencing still occurred without HUS1 and RAD9A (Lyndaker et al., 2013a; Vasileva et al., 2013). By contrast, the localization of ATR, TOPBP1 and BRCA1 to unsynapsed regions was compromised in *Rad1* CKO spermatocytes. ATR and BRCA1 work in a positive feedback loop to encourage meiotic silencing (Royo et al., 2013; Turner et al., 2004), and the canonical and alternative 9-1-1 complexes may also be part of this regulatory circuitry. Phosphorylation of some ATR targets, such as H2AX and HORMAD2, still occurred in *Rad1* CKO spermatocytes but only at a subset of unsynapsed chromatin regions. It should be noted that HORMAD1 and HORMAD2 localized appropriately to all unsynapsed regions independently of RAD1, indicating that HORMAD localization was not sufficient to driving ATR signaling and highlighting essential roles for the 9-1-1 complexes in meiotic ATR activation, possibly through interaction with TOPBP1. Other ATR substrates were more profoundly affected by RAD1 loss. CHK1 phosphorylation during meiosis was absent in *Rad1* CKO mice but present in *Hus1* CKO mice, suggesting that HUS1-independent alternative 9-1-1 complexes are necessary for meiotic CHK1 activation. CHK1 is required for timely ATR localization to the XY at mid-pachynema and MSCI initiation (Abe et al., 2018), and proper loading of RAD51 and DMC1 onto chromatin depend on CHK1 phosphorylation by ATR (Pacheco et al., 2018). Thus, the CHK1 phosphorylation defects described here when all 9-1-1 complexes are disrupted could contribute to multiple phenotypes observed in *Rad1* CKO spermatocytes, including impaired silencing and faulty DSB repair.

We also observed reduced SC protein phosphorylation in *Rad1* CKO and ATRi-treated mice, suggesting a role for the 9-1-1 complexes in mammalian SC formation. Studies in *S. cerevisiae* show that direct interaction between the 9-1-1 complex and an SC component, Red1, is required for both meiotic checkpoint signaling and SC formation (Eichinger and Jentsch, 2010). Additionally, the budding yeast 9-1-1 complex also directly interacts with Zip3, a member of the ZMM (Zip, Mer, Msh) group of proteins that promote initiation of SC formation and crossover recombination. Notably, budding yeast 9-1-1 and clamp loader mutants show reduced ZMM assembly on chromosomes, impaired SC formation, and reduced interhomolog recombination (Eichinger and Jentsch, 2010; Ho and Burgess, 2011; Shinohara et al., 2019; Shinohara et al., 2015).

ATR phosphorylates several cohesin complex components such as SMC1β and SMC3 (Fukuda et al., 2012). Phosphorylation of these proteins at the canonical ATR S/T-Q motif was downregulated in both ATRi-treated and *Rad1* CKO mice. SMC3 localization to meiotic chromosome cores was unperturbed, but SMC3 phosphorylation was dependent on RAD1 and ATR. It is important to note that additional proteins that are part of the cohesin complex, such as WAPL and SORORIN, also showed reduced phosphorylation in our phosphoproteomic screen. NIPBL, which functions in association with Mau2 as a SMC loader that localizes to chromosomal axes from zygonema to mid-pachynema (Visnes et al., 2014) also had reduced phosphorylation in testes from *Rad1* CKO and ATRi-treated mice. Interestingly, in *C. elegans*, SCC-2^NIPBL^ loss disrupts DSB processing, cohesin loading and 9-1-1 recruitment to DNA damage sites (Lightfoot et al., 2011).

In addition to phosphoproteomics, we used ERC analysis to reveal potential mechanistic roles for 9-1-1 subunits. ERC analysis can infer functional protein partners based upon correlated rates of evolutionary change. Consistent with our phosphoproteomics data, this analysis highlighted significant evolutionary correlations between the genes encoding 9-1-1 complex subunits and those encoding proteins involved in SC formation, such as SYCP1, SYCE1, SYCE1L and SYCE2, in addition to RAD21, RAD21L and SMC1β which are involved in cohesion. Defects in homolog synapsis in *Rad1* CKO mice, together with the decreased cohesin phosphorylation, further implicates the 9-1-1 complexes in these key aspects of meiotic chromosome structure. However, further exploration of the mechanisms underlying the interactions between SC proteins, cohesin, and the 9-1-1 complexes is necessary and may provide insights into the basis for the DSB repair defects in *Rad1* CKO mice, as proper SC formation and cohesin function is important for DSB repair (Ishiguro, 2019).

In mitotic cells, ATR activation is dependent on the 9-1-1/TOPBP1 axis under cellular stress, while ATR activation during unperturbed conditions relies on ETAA1 (Bass and Cortez, 2019). The potential contributions of ETAA1 to meiotic ATR activation have yet to be directly assessed. Our phosphoproteomic screen showed that RAD1-independent, ATR-dependent differentially phosphorylated proteins were associated with top gene ontology terms of cellular processes and organelle organization (Sims et al., 2021). Mice expressing a ETAA1 mutant with a 42 amino acid deletion show signs of replication stress but are fertile (Miosge et al., 2017). Understanding the differential roles of 9-1-1/TOPBP1 and ETAA1 in meiotic ATR activation may highlight different modes of structure-specific ATR activation that are coupled with distinct downstream outputs.

Although this study highlights key meiotic functions of both canonical and alternative 9-1-1 complexes, our approach does not resolve the relative importance of the DNA repair and checkpoint signaling roles of the 9-1-1 complexes during meiosis. Previous studies identified separable roles for 9-1-1 complexes in ATR activation via TOPBP1 interaction, and DNA repair protein scaffolding through the outer surface of 9-1-1 clamps (Lim et al., 2015). The loss of 9-1-1 complex formation and loading in *Rad1* CKO mice disrupts both of these roles. In budding yeast, the direct interactions between the 9-1-1 complex and Red1 as well as Zip3, together with additional evidence that the roles for 9-1-1 in SC formation and recombination can be distinguished from those of Mec1 (ATR), provide compelling support for the notion that the 9-1-1 complex executes signaling-independent functions during meiosis, aside from its roles in checkpoint signaling (Eichinger and Jentsch, 2010; Shinohara et al., 2019; Shinohara et al., 2015). In the future, separation-of-function 9-1-1 mouse mutants could be used to clarify precisely how the 9-1-1 complexes mediate meiotic processes such as homolog synapsis, cohesion, and silencing. Moreover, continued genetic and biochemical analysis of the paralogs RAD9B and HUS1B holds promise for resolving the differential and overlapping roles of the canonical and alternative 9-1-1 complexes in spermatogenesis.

## MATERIALS AND METHODS

### Mice and genotyping

*Rad1* CKO and control mice on the 129Sv/Ev background were generated by crossing *Rad1^flox/flox^* mice with *Rad1^+/+^, Stra8-Cre^+^* mice to generate *Rad1^+/fl^, Stra8-Cre^+^ (Rad1^+/-^, Stra8-Cre^+^)* mice. *Stra8-Cre* mice containing one null *Rad1* allele (*Rad1^+/-^, Stra8-Cre^+^)* were crossed with *Rad1^flox/flox^* mice to generate experimental germ-cell specific *Rad1* conditional knockout mice (*Rad1^-/fl^, Stra8-Cre^+^)* and control mice (*Rad1^+/fl^, Stra8-Cre^+^; Rad1^+/fl^, Stra8-Cre^-^; Rad1^-/fl^, Stra8-Cre^-^*). *Rad1^flox^* mice carry a conditional *Rad1* allele containing a K185R mutation that does not affect RAD1 function (Wit et al., 2011). *Hus1* conditional knockout mice were used as previously reported (Lyndaker et al., 2013a). All mice used for this study were handled following federal and institutional guidelines under a protocol approved by the Institutional Animal Care and Use Committee (IACUC) at Cornell University. The key resources table lists the genotyping primers used in this study.

### Fertility tests

For fertility testing, 8-to 12-week-old *Rad1^-/fl^, Stra8-Cre^+^* and control males were singly housed with wild-type FVB females, where copulatory plugs were monitored daily. Once a plugged female was detected the female was removed to a separate cage and monitored for pregnancy. Viable pups were counted on the first day of life.

### Epididymal sperm counts

Both caudal epididymides from 12-week-old mice were minced with fine forceps in 37°C in a petri dish containing 1x Phosphate Buffered Saline (PBS) and fixed in 10% neutral-buffered formalin (1:25 dilution). Sperm were counted using a hemacytometer and analyzed statistically using a Student’s t-test between control and *Rad1* CKO mice.

### Irradiation of mice

Control and *Rad1* CKO mice were placed in a ^137^Cesium sealed source irradiator (J.L. Shepherd and Associates) with a rotating turntable and irradiated with 5Gy IR. Testes were harvested for meiotic spreads 1-hour post radiation.

### Immunoblotting

Whole testis lysates from *Rad1* CKO, *Hus1* CKO, and control mice were prepared in RIPA buffer (10mM Tris-HCl, pH 8.0, 1mM EDTA, 0.5mM EGTA, 1% Triton X-100, 0.1% Sodium Deoxycholate, 0.1% SDS, 140mM NaCl) supplemented with aprotinin, leupeptin, sodium orthovanadate, and phenyl-methylsulfonyl fluoride. Cell lysates were resolved by SDS-PAGE and immunoblotted using standard procedures. Bands were visualized on a VersaDoc MP 5000 Model (Bio-Rad) using a 1:1 ratio of WesternBright ECL Luminol/enhancer solution to WesternBright Peroxide Chemiluminescent peroxide solution (Advansta). Antibody information is provided in key resources table.

### Histology and immunohistochemistry

Testes were harvested from mice aged to 8dpp, 4 weeks or 12 weeks of age. Testes were then fixed overnight in either Bouin’s (Ricca chemicals) for hematoxylin and eosin staining or 10% neutral-buffered formalin (Fisher) for LIN28, TRA98, and TUNEL staining. Fixed testes were embedded in paraffin wax and sectioned at 5µm. Immunofluorescence staining was used to detect LIN28 using rabbit polyclonal anti-LIN28 antibody (Abcam, ab63740). Immunohistochemistry staining was used to detect TRA98 using rat monoclonal anti-TRA98 antibody (BioAcademia, 73-003). TUNEL assay was performed using the Apoptag^®^ kit (EMD Millipore) as per the manufacturer’s instructions. LIN28, TRA98 and TUNEL data were quantified in ImageJ by counting the number of positive cells per tubule for 50 tubules of each genotype for each age group. Differences between controls and *Rad1* CKOs was analyzed using Welch’s unpaired t-test using Graphpad.

### Meiotic spreading and immunofluorescence staining

Meiotic spreads were prepared from 8-to 12-week-old mice as previously described (Kolas et al., 2005). Briefly, tubules from mice were incubated on ice in hypotonic extraction buffer for 1 hour. Tubules were than minced into single cell suspension in 100mM sucrose, and cells were spread on slides coated with 1% PFA with 0.15% TritionX-100 and incubated in a humidifying chamber for 4 hours or overnight. For immunostaining, slides were blocked using 10% goat serum and 3% BSA, followed by incubation overnight with primary antibody (listed in key resources table) at room temperature in a humidifying chamber. Secondary antibodies were incubated at 37°C for 2 hours in the dark, and slides were then cover-slipped using anti-fade mounting medium (2.3% DABCO, 20mM Tris pH 8.0, 8µg DAPI in 90% glycerol). Meiotic chromosomal spreads were imaged with an AxioCam MRM using a Zeiss Imager Z1 microscope (Carl Zeiss, Inc.) and processed with ZEN Software (version 2.0.0.0;Carl Zeiss, Inc.). Quantification of meiotic spreads was performed using Fiji for ImageJ. Statistical analysis was performed using Welch’s unpaired t-test using Graphpad Prism8.

### RNA Fluorescence In-Situ Hybridization (RNA-FISH) and immunofluorescence Staining

RNA FISH was carried out with digoxigenin labelled probe using BAC DNA, *Scml2*: RP24-204O18 (CHORI) and immunofluorescence using rabbit HORMAD2, antibody (gift from gift from A. Toth) as previously described (Mahadevaiah et al., 2009). Images of RNA FISH with immunofluorescence were captured using Deltavision Microscopy System (100x/1.35NA Olympus UPlanApo oil immersion objective.

### ATR inhibitor treatment of mice

Wild-type B6 mice were treated via oral gavage with AZ20 (Selleck Chemicals) reconstituted in 10% DMSO (Sigma), 40% Propylene glycol (Sigma), and 50% water. Three different ATRi treatments were used. Chronic ATR inhibitor treatment was performed by treating mice daily for 3 days with 50mg/kg AZ20 and collecting 24 hours after final dose, or by treating with 2 doses of 50mg/kg AZ20 on the first 2 days and 1 final dose of 25mg/kg AZ20 and collecting 4 hours after the final dose. Acute ATR inhibitor-treated mice were collected 4 hours after one dose of 50mg/kg AZ20.

### Phosphoproteomic analysis

Whole testis lysates from 8-week-old AZ20 (ATRi) treated wild-type C57BL/6 and 12-week-old *Rad1* CKO mice were subjected to phosphopeptide enrichment and 6-plex TMT (ThermoFisher) labeling (Sims et al., 2021). To identify differentially regulated phosphosites we performed a bow tie analysis as described in Sims et al. (Sims et al., 2021). Gene ontology enrichment with Benjamini-Hochberg adjustment to account for multiple hypothesis testing was done using STRINGdb. Terms were ranked based on their FDR values, and the top 10 gene ontology terms from the Biological Processes subontology were plotted using R package GOplot.

### Orthology analysis

Human 9-1-1 subunit sequences were used to obtain their respective orthologs from Ensemble 101(2020) and/or NCBI Gene from 33 representative mammalian species. Orthologs found in Ensemble having a ≥50% of both target and query sequence identity and a pairwise whole genome alignment score of ≥50 were considered to have high confidence. Orthologs that did not meet those criteria were considered to have low confidence. Sequences only found in NCBI Gene database were considered as high confidence if they were found to be syntenic. Synteny was determined based on whether the gene had at least one shared neighbor gene upstream or downstream that also was conserved. Species divergence across time was obtained from TimeTree website (www.timetree.org).

### Phylogenetic analysis

Protein sequences of 9-1-1 orthologs were obtained using NCBI HomoloGene. Multiple alignment of protein sequences was done using Clustal Omega (1.2.2) implemented in Geneious Prime (2020.0.5). A substitution model was tested using ProtTest (v. 3.4.2). The selected substitution model with specific improvements was JTT+I+G+F (Jones-Taylor-Thornton; +I: invariable sites; +G: rate heterogeneity among sites; +F: observed amino acid frequencies). Improvements were included to take account for any evolutionary limitations due to conservation of protein structure and function. A nonrooted phylogenetic tree was made using Maximum Likelihood interference (4 gamma distributed rate) (Nguyen et al., 2015) and implemented with iTOL (itol.embl.de) (Letunic and Bork, 2019). Branch distance represents substitution rate and branch support was performed with 1000 ultrafast bootstrap replicates. Nodes below 70% branch support were collapsed.

### ERC analysis

ERC calculations were completed using the Evolutionary Rate Covariation (ERC) web tool at https://csb.pitt.edu/erc_analysis/ (Wolfe and Clark, 2015). Group analysis was performed to examine ERC values between all gene pairs indicated in Figure 2A, 7A and 7B using UCSC gene sequences from 33 mammalian species as described in Priedigkeit et al. (Priedigkeit et al., 2015). For figure 7-figure supplement 1A and 1B, the protein set list for Gene Ontology subontology Meiosis I (GO:0007127) was obtained from AmiGO 2 (v2.5.13). ERC values were calculated against each of the 9-1-1 subunits using the ERC analysis web site. Using R (v4.0.3) ERC values were depicted as a heatmap and a network plotted using the packages pheatmap (v1.0.12) and qgraph (v1.6.5) respectively. A cutoff of ERC value of 0.4 was used to determine significant comparisons. The Fruchterman & Reingold algorithm was used to generate a forced-directed layout to help determine clusters of highly connected nodes and after 500 iterations the distance between nodes shows absolute edge weight (ERC values) between nodes.

## ACKNOWLEDGMENTS

We are thankful to Dan Barbash and Eric Alani for helpful discussions and for providing critical feedback on the manuscript, to Mary Ann Handel and Attila Toth for providing reagents used in this study, and to Christina Jeon for early-stage contributions to the analysis of 9-1-1 subunit evolution. This work was supported in part by NIH grants R03 HD083621 (to RSW), R01 HD095296 (to MBS and RSW), R01 HD097987 (to PEC), NSF predoctoral fellowship DGE-1144153 (to CP), and a National Center for Research Resources instrumentation grant (S10 RR023781). This work additionally was supported by European Research Council (CoG 647971) and the Francis Crick Institute, which receives its core funding from Cancer Research UK (FC001193), UK Medical Research Council (FC001193) and Wellcome Trust (FC001193).

## AUTHOR CONTRIBUTIONS

Conceptualization: CP, AML, MAB-E, PEC, MBS, RSW; Methodology and experimentation: CP, GAAM, MAB-E, MG, MSD, EK, KJG, SKM, CT, CJS, JS; Data curation and analysis: CP, GAAM, VMF, MG, MSD, SKM, CT, CJS, JS; Project administration and funding: JMAT, PEC, MBS, RSW; Resources: NW, HJ, NLC, PEC, RF; Supervision: JMAT, PEC, MBS, RSW; Writing and editing: CP, GAAM, AML, JS, MAB-E, PEC, MBS, RSW.

## KEY RESOURCES TABLE

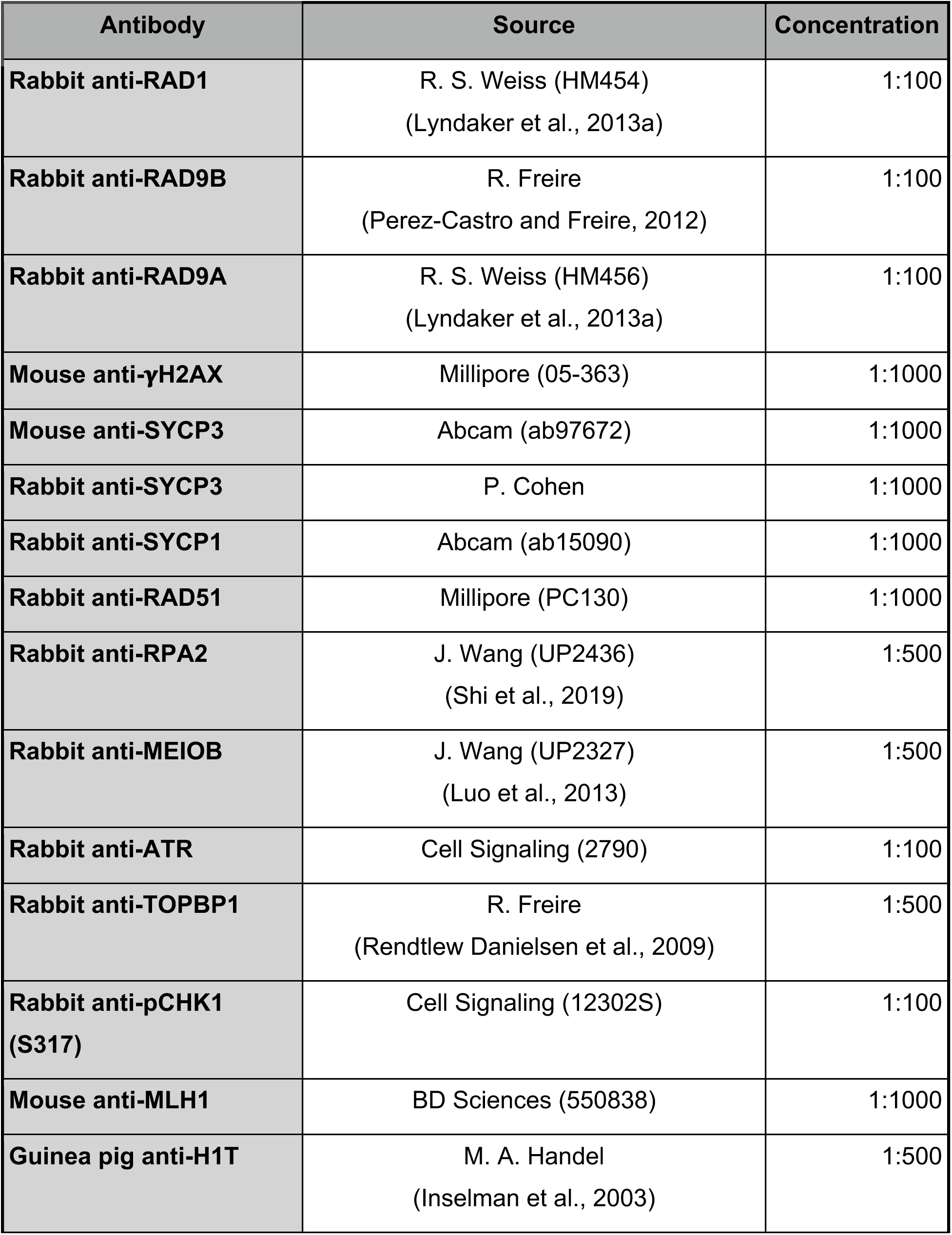
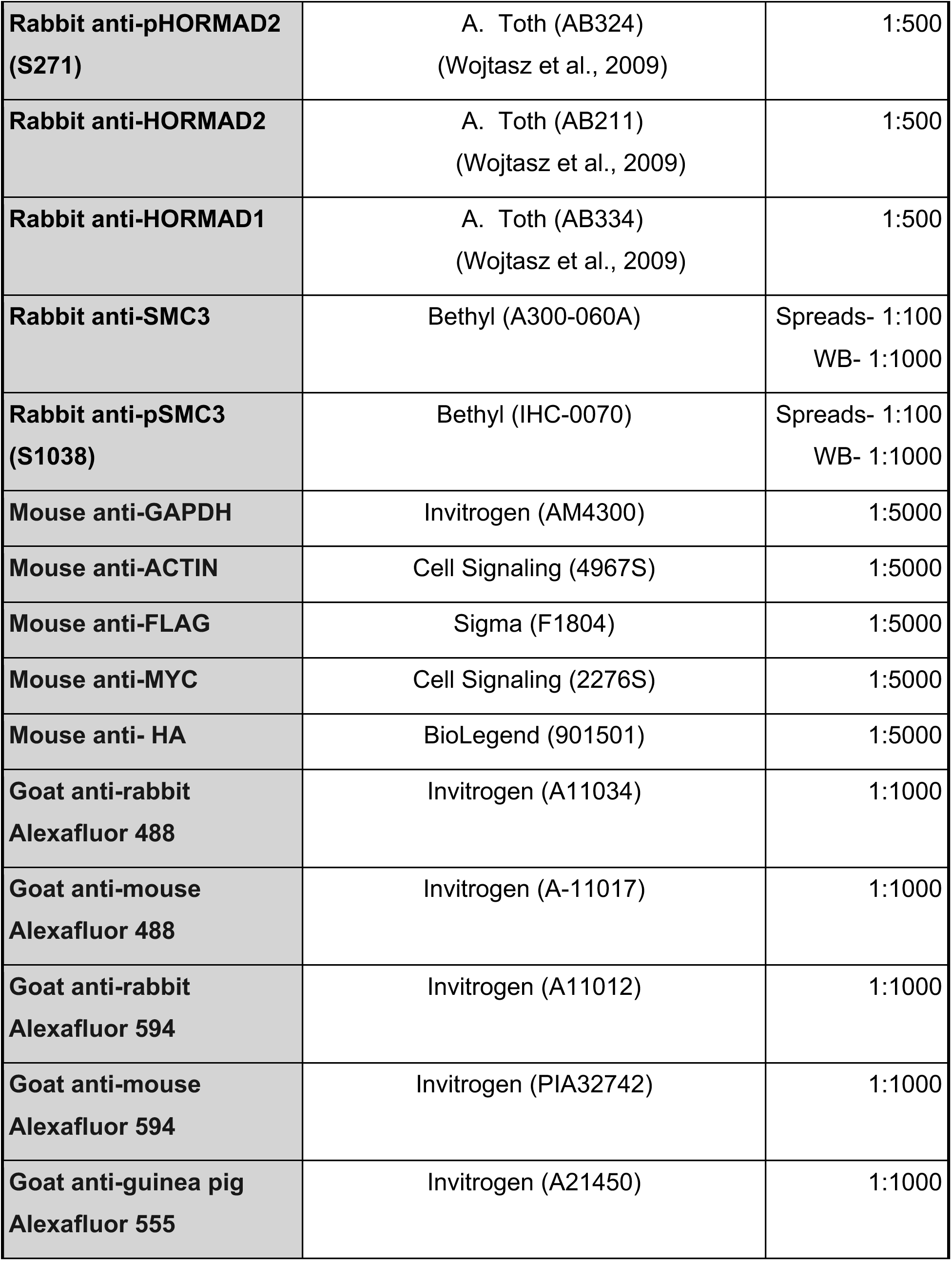
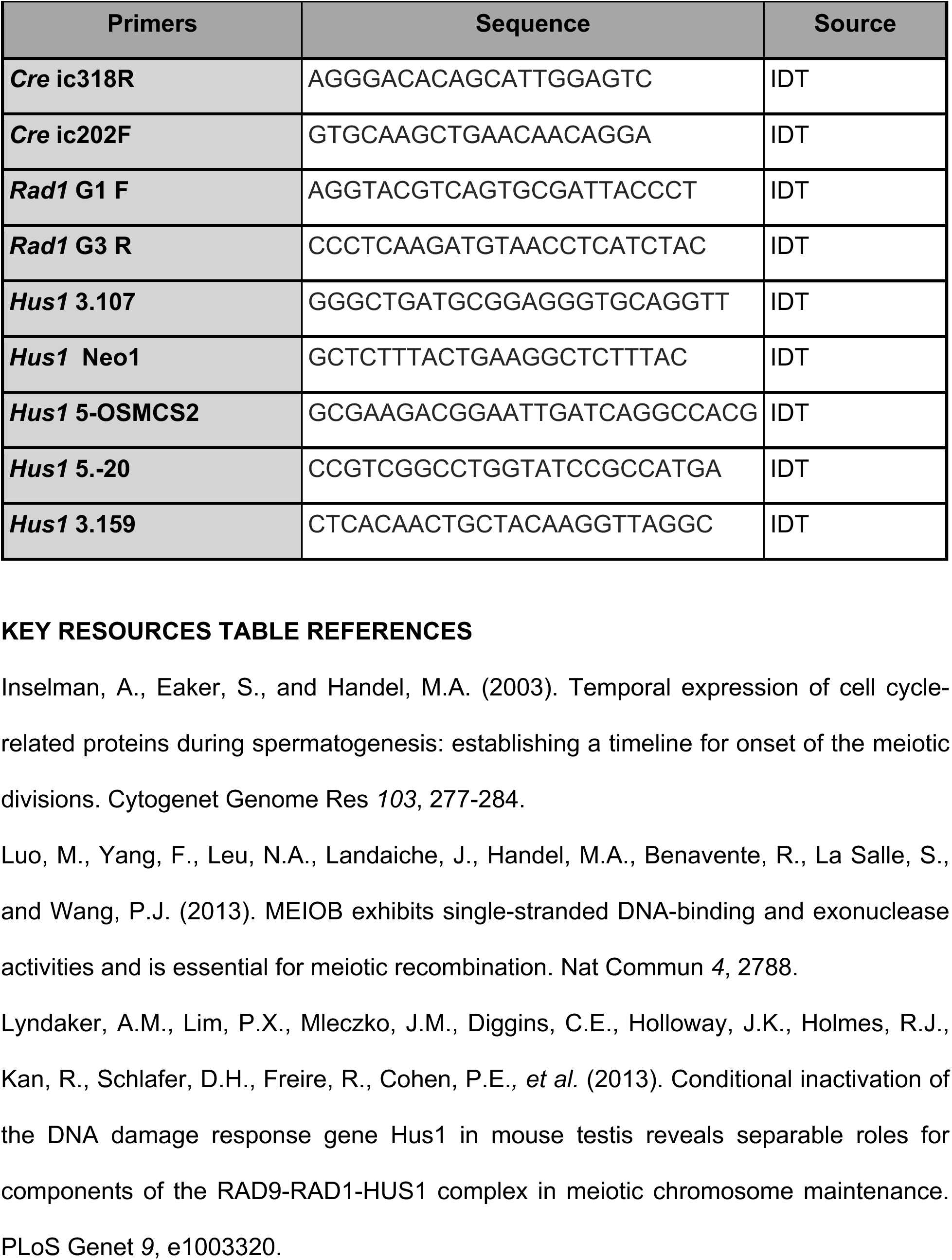
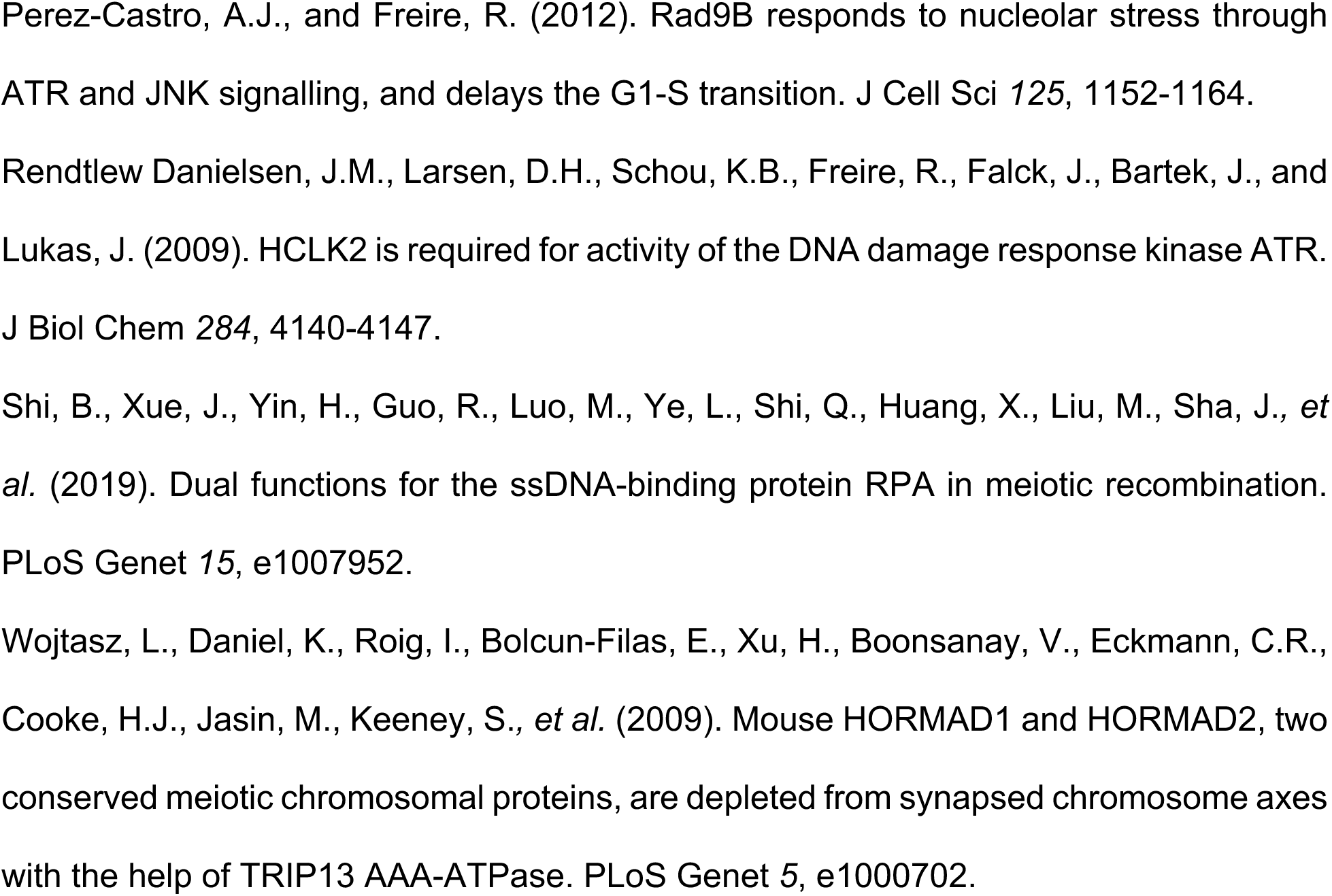

---

**Figure 1 Supplement 1:**
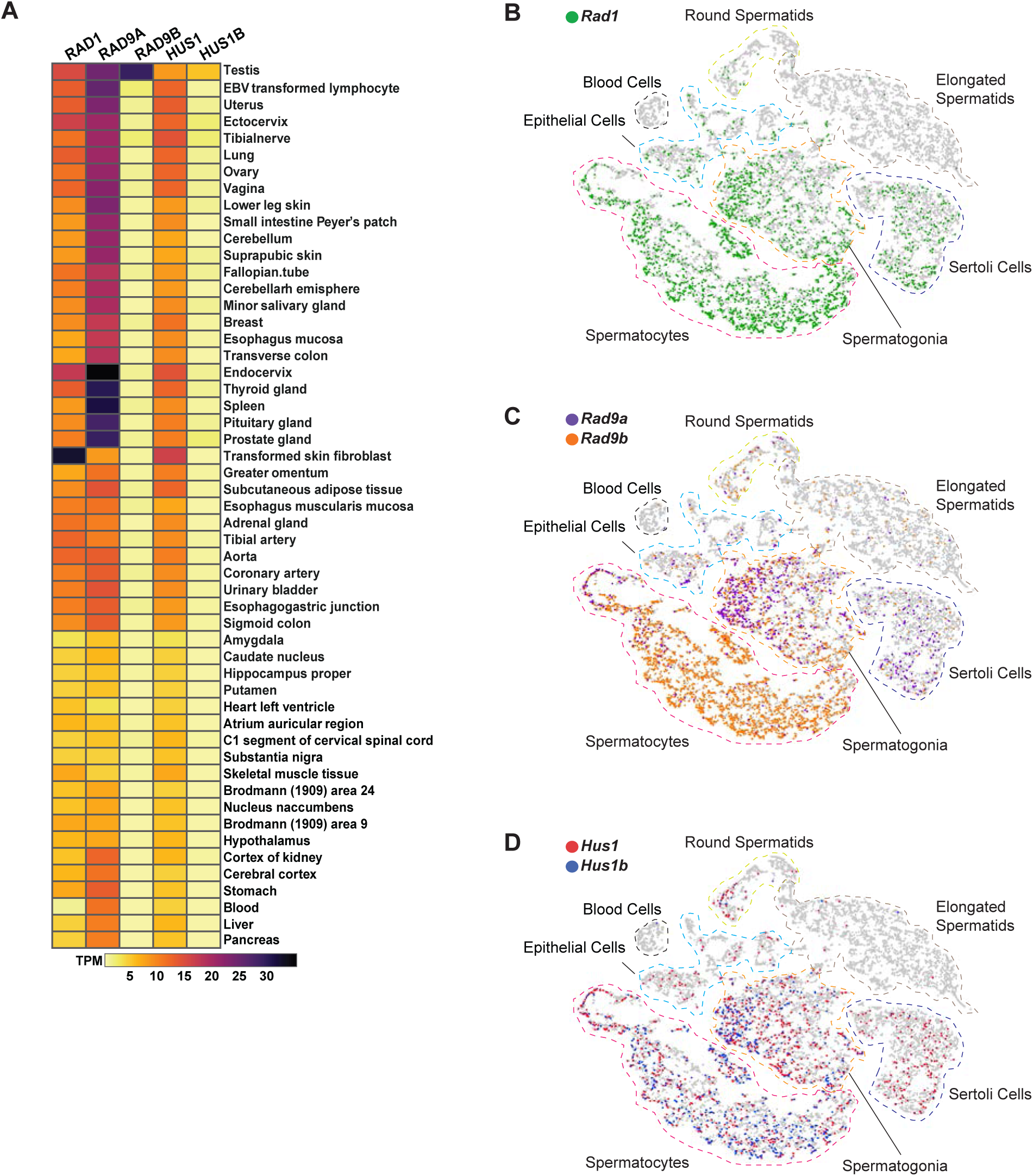
Expression of 9-1-1 complex subunits. (A) Expression of 9-1-1 subunits in various human tissues. Data from the Genotype-Tissue Expression (GTEx) project was obtained in Expression Atlas – EMBL-EBI. Gene expression values are shown as transcript per million (TPM). (B-D) tSNE plots of single-cell RNA seq analysis of mouse testes demonstrating the expression of 9-1-1 subunits in single cells from round spermatids, elongated spermatids, blood cells, epithelial, spermatocytes, spermatogonia, and Sertoli cells population within testes. Gray circles are individual cells. *Rad1*-expressing cells are shown in green (B). *Rad9a*-expressing cells are shown in purple, and *Rad9b-*expressing cells are in orange (C). *Hus1*-expressing cells are in red, and *Hus1b*-expressing cells are in blue (D).

**Figure 2 Supplement 1:**
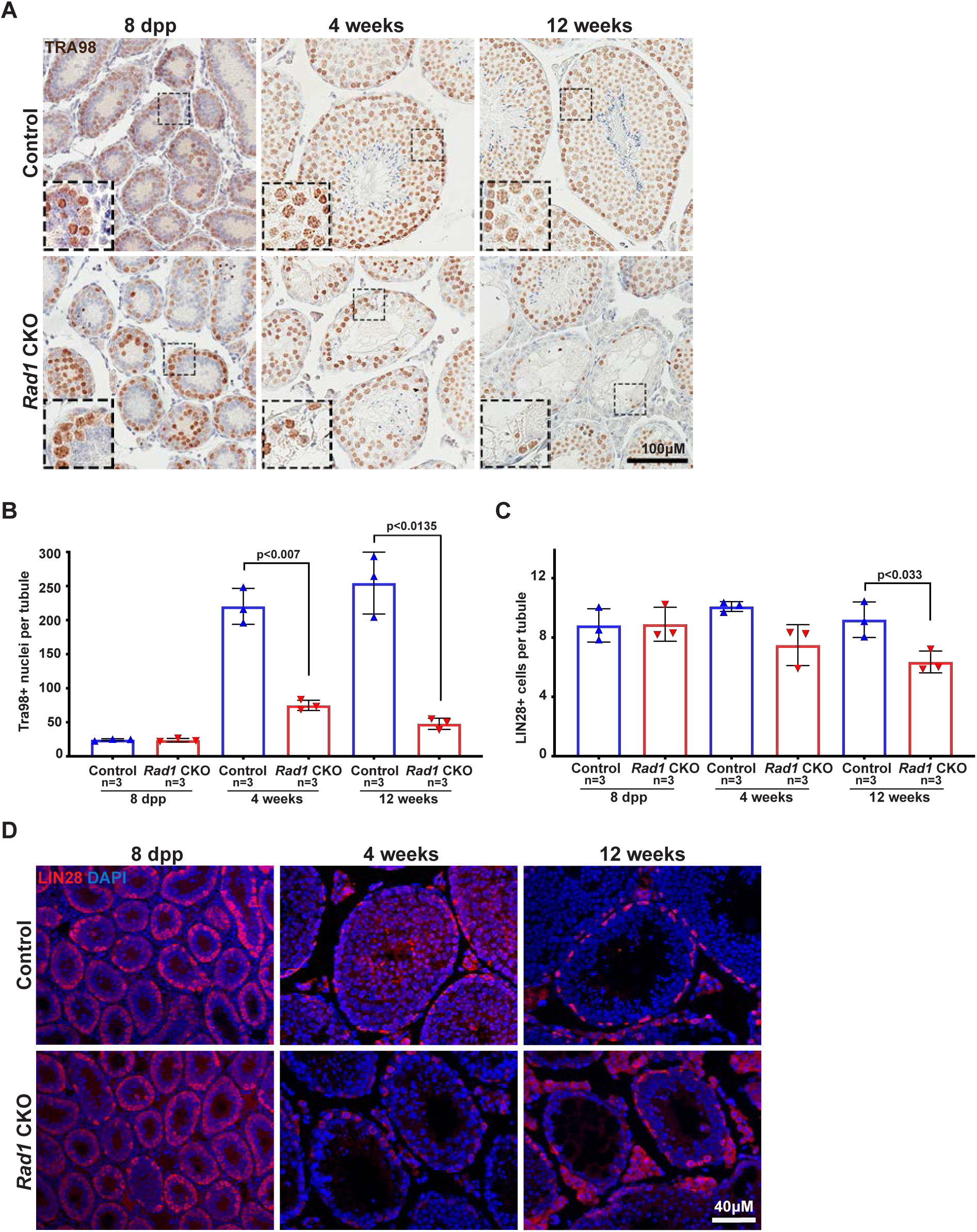
*Rad1* inactivation in testis causes germ cell loss. (A-B) Representative images (A) and quantification (B) of TRA98-positive cells per tubule in control and *Rad1* CKO mice (n= number of mice; 50 tubules per mouse quantified). (C-D) Representative images (C) and quantification (D) of LIN28-positive spermatogonial stem cells (n= number of mice; 50 tubules per mouse quantified). p-value calculated using Welch’s unpaired t-test in Graphpad Prism8.

**Figure 3 Supplement 1:**
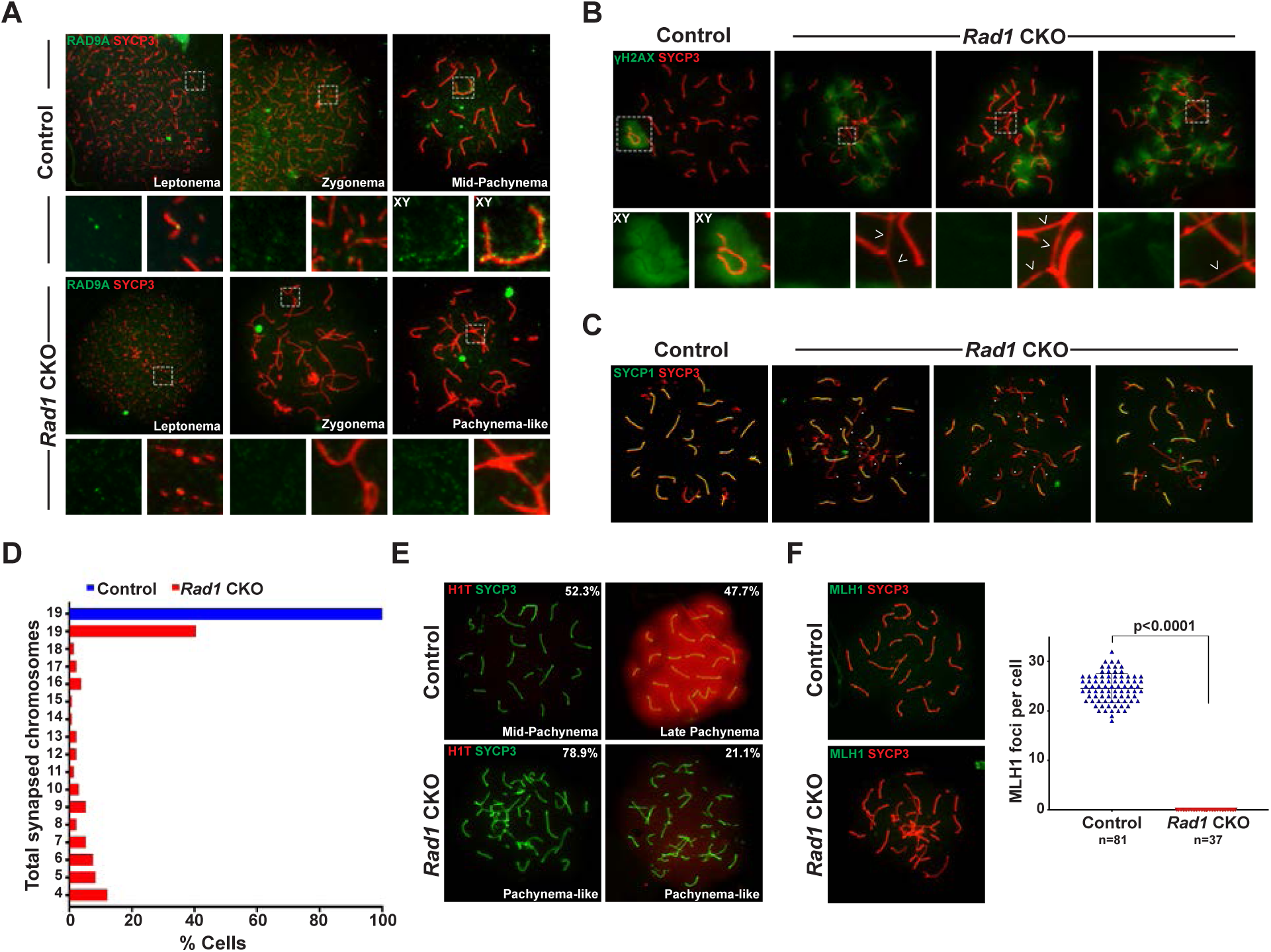
*Rad1* CKO spermatocytes show RAD9A loss and do not progress to mid-pachynema. (A) Representative meiotic spread images stained for RAD9A from control and *Rad1* CKO 12-week-old mice (3 control mice; n= 178 cells; 3 CKO mice; n=81 cells). (B) Additional examples of yH2AX meiotic spread staining in control and *Rad1* CKO mice (3 control mice; n= 127 cells; 5 CKO mice; n=205 cells). (C) Examples of SYCP1/3 co-staining in control and *Rad* CKO meiotic spreads, asterisks indicate properly synapsed chromosomes (3 control mice; n= 156 cells; 3 CKO mice; n=131 cells). (D) Total synapsed chromosomes per cell in control (blue) and *Rad1* CKO (red) per cell (3 control mice; n= 156 cells; 3 CKO mice; n=131 cells). (E) H1T meiotic spread staining from control and *Rad1* CKO mice. Of 174 pachynema staged cells analyzed from control mice 52.3% cells showed no H1T staining (2 control mice). 104 pachynema-like staged cells from *Rad1* CKO mice were analyzed with 78.9% of cells displaying no H1T and 21.1% showed low levels of H1T staining (2 CKO mice). (F) Representative images of MLH1 staining in control and *Rad1* CKO spreads, and quantification of MLH1 foci (D) (2 control mice; n= 81 cells; 2 *Rad1* CKO mice; n=37 cells). p-value calculated using Welch’s unpaired t-test using Graphpad Prism8.

**Figure 4 Supplement 1:**
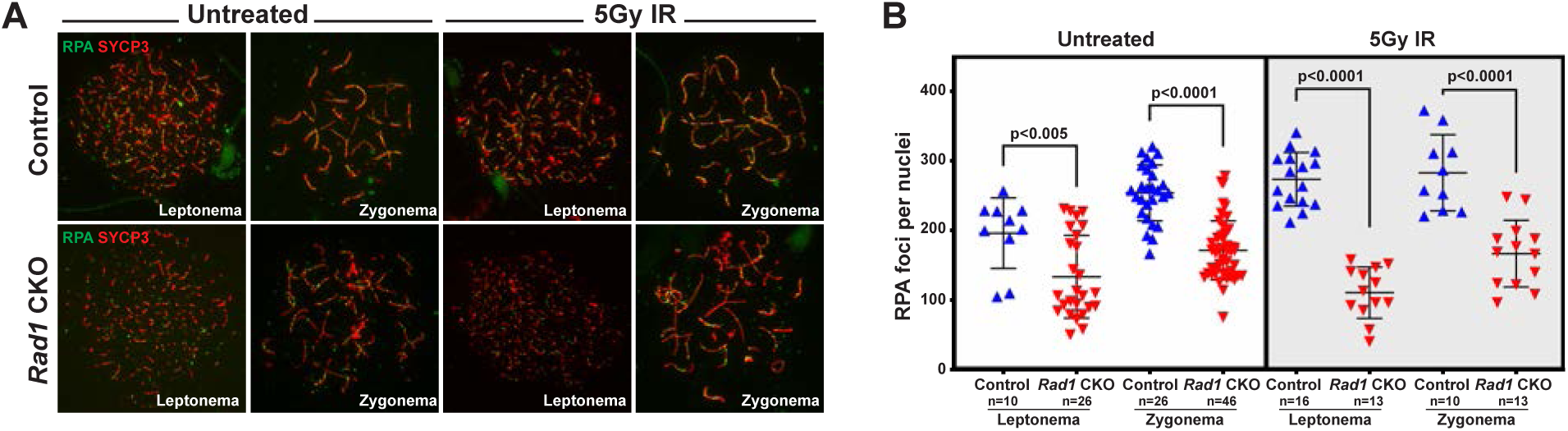
Repair of IR-induced DSBs during early prophase I is compromised in the absence of 9-1-1 complexes. 8-week-old control and *Rad1* CKO mice were irradiated with 5Gy IR and collected 1-hour post IR. (A and B) Representative images (A) and quantification (B) of RPA staining of meiotic spreads prepared from mice of the indicated genotypes (2 control mice and 2 CKO analyzed; n=total cells analyzed). (C and D) MEIOB representative meiotic spreads images (C) and quantifications (D) (2 control mice and 2 CKO analyzed; n=total cells analyzed). p-value calculated using Welch’s unpaired t-test using Graphpad Prism8.

**Figure 7 Supplement 1:**
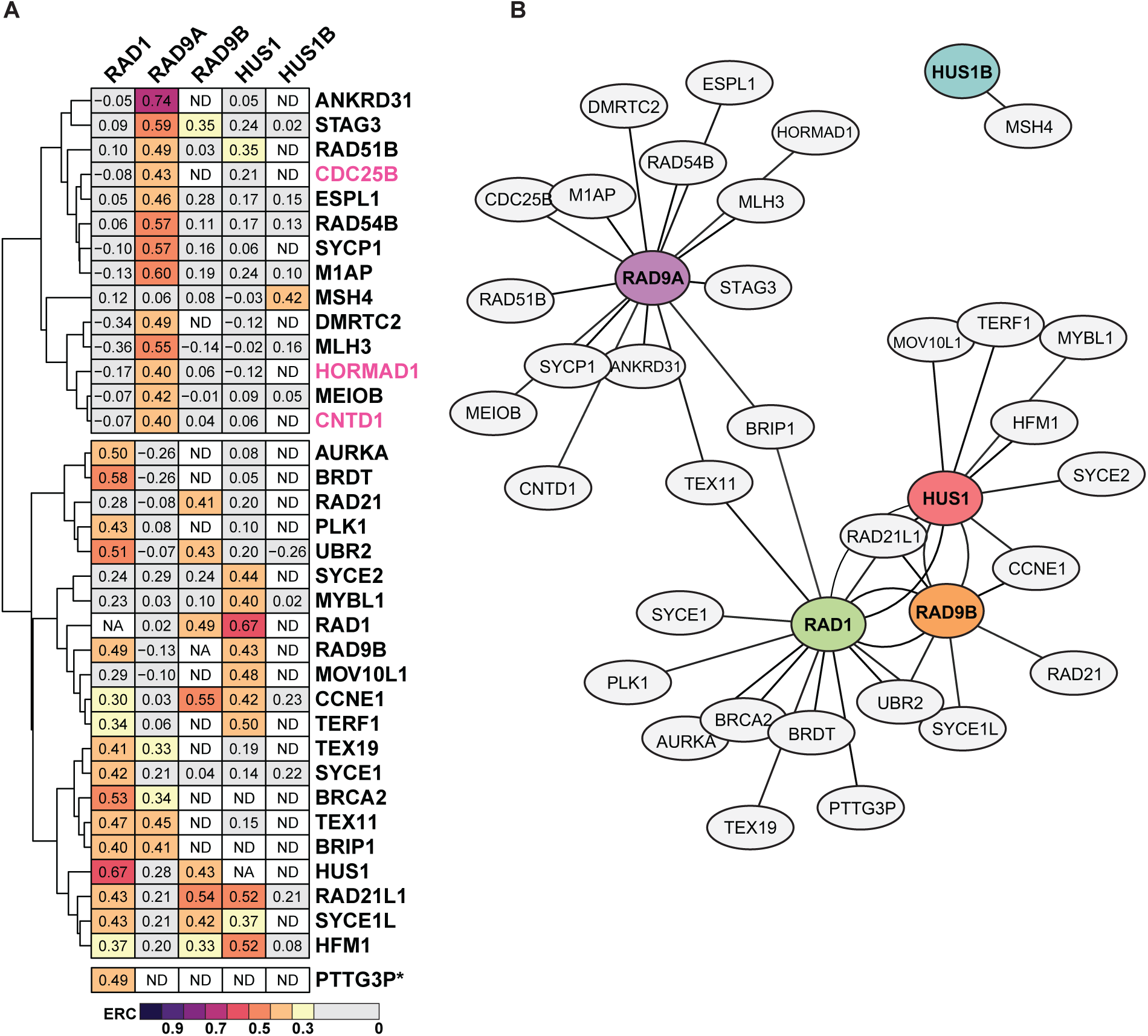
Evolutionary rate covariation network of meiosis I proteins. (A) Heatmap of proteins found under Gene ontology term meiosis I (GO:0007127) that showed ERC values ≥0.4 for each subunit. Pink labeling denotes proteins with no significant ERC values (p>0.05). ND: no value exists in dataset for paired comparison between the two proteins. NA: not applicable for comparisons between the same protein. PTTG3P* has a high ERC and significance with RAD1 only. (B) ERC values between subunits and meiosis I proteins were used to plot the network with force-directed layout. The Fruchterman & Reingold algorithm features attraction between highly connected nodes, identifying protein clusters based on ERC data. The distance between nodes is proportional to absolute edge weight (ERC value of each protein), with shorter distances between nodes reflecting higher ERC values.

